# Spatially resolved single-cell multiomics map of human trophoblast differentiation in early pregnancy

**DOI:** 10.1101/2022.11.06.515326

**Authors:** Anna Arutyunyan, Kenny Roberts, Megan A Sheridan, Ilia Kats, Luz Garcia-Alonso, Britta Velten, Regina Hoo, Kevin Troulé Lozano, Louis-Francois Handfield, Luca Marconato, Elizabeth Tuck, Lucy Gardner, Cecilia Icoresi Mazzeo, Iva Kelava, Elena Prigmore, Sarah A Teichmann, Omer Ali Bayraktar, Ashley Moffett, Oliver Stegle, Margherita Y Turco, Roser Vento-Tormo

**Author notes:** co-first (Anna Arutyunyan, Kenny Roberts, Megan A Sheridan). co-last and co-corresponding (,).

## Abstract

The relationship between the human placenta, the extraembryonic organ built by the fetus, and the decidua, the mucosal layer of the uterus, is essential to nurture and protect the fetus during pregnancy. Extravillous trophoblast cells (EVTs) anchor the placenta and infiltrate the decidua, transforming the maternal arteries into high conductance vessels. Defects in trophoblast invasion and arterial transformation established during early pregnancy underlie common pregnancy disorders such as pre-eclampsia. Despite its importance, how EVT invasion is regulated in humans is still unclear due the inaccessibility of the entire pregnant uterus and, until recently, a lack of reliable *in vitro* models. Here, we have generated a spatially-resolved multiomics single-cell atlas of the entire maternal-fetal interface including the myometrium, allowing us to resolve the full trajectory of trophoblast differentiation. We have used this cellular map to elucidate the main regulatory programmes mediating EVT invasion and show that they are preserved in trophoblast organoids. We define the transcriptomes of the final cell states of trophoblast invasion: placental bed giant cells (fused multinucleated EVTs) and endovascular EVTs (which form plugs inside the maternal arteries). We reconstruct the cell-cell communication events contributing to trophoblast invasion and GC formation, and define the dual role of interstitial EVTs and endovascular EVTs in mediating arterial transformation during early pregnancy. Together, our data provides a comprehensive analysis of postimplantation trophoblast differentiation in humans that can be used as a blueprint to design accurate multilineage placental *in vitro* models.

---

During the nine months of human pregnancy the fetus is entirely dependent on its placenta. This transient extra-embryonic organ is located at the interface between the mother and her fetus. Trophoblast is the main cell type of the placenta, and arises from the trophectoderm surrounding the preimplantation embryo^1^. After implantation, extravillous trophoblast cells (EVTs) emerge from the cytotrophoblast shell, infiltrate the decidua, the mucosal layer of the pregnant uterus, and migrate towards the spiral arteries where they destroy the smooth muscle media. Subsequently, endovascular trophoblast cells (eEVTs) form a plug close to the cytotrophoblast shell where the arteries terminate and replace the endothelium^2^. In this way EVTs transform maternal arteries in the decidua basalis into high conductance vessels^3–6^. EVTs begin to fuse into placental bed giant cells (GCs) deeper in the decidua and eventually migrate as far as the inner third of the myometrium^7^.

Defects in decidualisation are associated with pre-eclampsia^8^, a syndrome characterised by defective arterial transformation by EVTs. In contrast, excessive invasion of EVTs into the uterus occurs when the decidua is missing (for instance at a scar from a previous caesarean section) and can even cause uterine rupture^9^. Thus, placentation and successful pregnancy depends on the correct degree of trophoblast invasion, and the decidua plays an important role. Both trophoblast cell-intrinsic mechanisms (i.e. precisely coordinated gene expression as EVTs invade) and signals provided by the surrounding maternal decidual cells contribute to this crucial process.

Investigating the human maternal-fetal interface early in pregnancy is hampered by ethical and logistical limitations because samples can only be obtained from voluntary terminations of pregnancy. Moreover, animal models are of limited use in modelling the particularly invasive haemochorial type of placentation characteristic of humans, which is distinct even from other primates apart from great apes^10^. Primary trophoblast organoids are able to recapitulate some aspects of placental development and invasion^11–13^ but their accuracy at the single-cell level remains to be determined. Our previous single-cell transcriptomics analysis of the first trimester maternal-fetal interface has provided an unprecedented view of the cell states comprising this environment^14^. However, the full spectrum of trophoblast states is not likely to be captured in existing single-cell transcriptomics atlases^14,15^ due to the absence of certain trophoblast subsets from decidual and placental tissue cell isolates. In particular, trophoblast cells present in the deeper layers of the decidua and myometrium are absent from standard surgical samples, and the villous syncytiotrophoblast (SCT), a multinucleated layer, is lost in classical single-cell RNA sequencing (scRNA-seq). A further difficulty is the loss of spatial context in these samples, which is essential to systematically resolve the interactions between trophoblast and decidual cells in early pregnancy.

Single-cell and spatial transcriptomic atlases of tissues have been transformative in understanding human development^16–19^, mapping disease^20,21^ and engineering organoids^22,23^. Here, we present a spatially-resolved single-cell multiomics characterization of the maternal-fetal interface. We examine the site of placentation from historical samples of first trimester hysterectomies, which include the entire uterus containing the placenta, decidua and myometrium. To faithfully recapitulate the dynamics of trophoblast invasion, we developed StOrder, a computational and statistical framework that reconstructs the smooth transition of cell states in space. Spatiotemporal ordering of trophoblast invasion allows us to characterise the molecular processes underpinning trophoblast invasion. We use this comprehensive detailed account of trophoblast differentiation to benchmark our trophoblast organoid model^11^.

Using our tool CellPhoneDB v4^18^, we describe interactions between trophoblast subsets and decidual cells that are likely to affect how trophoblast transformation of arteries occurs in early pregnancy. Thus, we provide a description of the whole trajectory of human trophoblast cell states and their spatial niches.

## Spatiotemporal map of the placental-decidual interface defines four villous cytotrophoblast subsets

We profiled three human implantation sites (between 6 and 9 post-conceptional weeks, PCW) using a multimodal approach **(Fig. 1a-c, Supplementary Tables 1-3)**. Consecutive sections from frozen tissue blocks of the implantation site were used for: (i) single-nuclei RNA sequencing (snRNA-seq); (ii) combined single nuclei RNA and ATAC sequencing (snRNA-seq/snATAC-seq, further referred to as multiome); and (iii) spatial transcriptomics using Visium. To account for the large tissue area of one donor (P13), we targeted four consecutive sections with four spatial transcriptomics capture areas **(Supplementary Fig. 1a)**. We also profiled five decidual and three placental samples from 8-13 PCW by scRNA-seq/snRNA-seq and integrated all the data with our previous scRNA-seq dataset of the maternal-fetal interface^14^ **(Fig. 1d, Supplementary Fig. 2a-e).** Our single-cell and spatial transcriptomics map is available at reproductivecellatlas.org.

**Figure 1.**
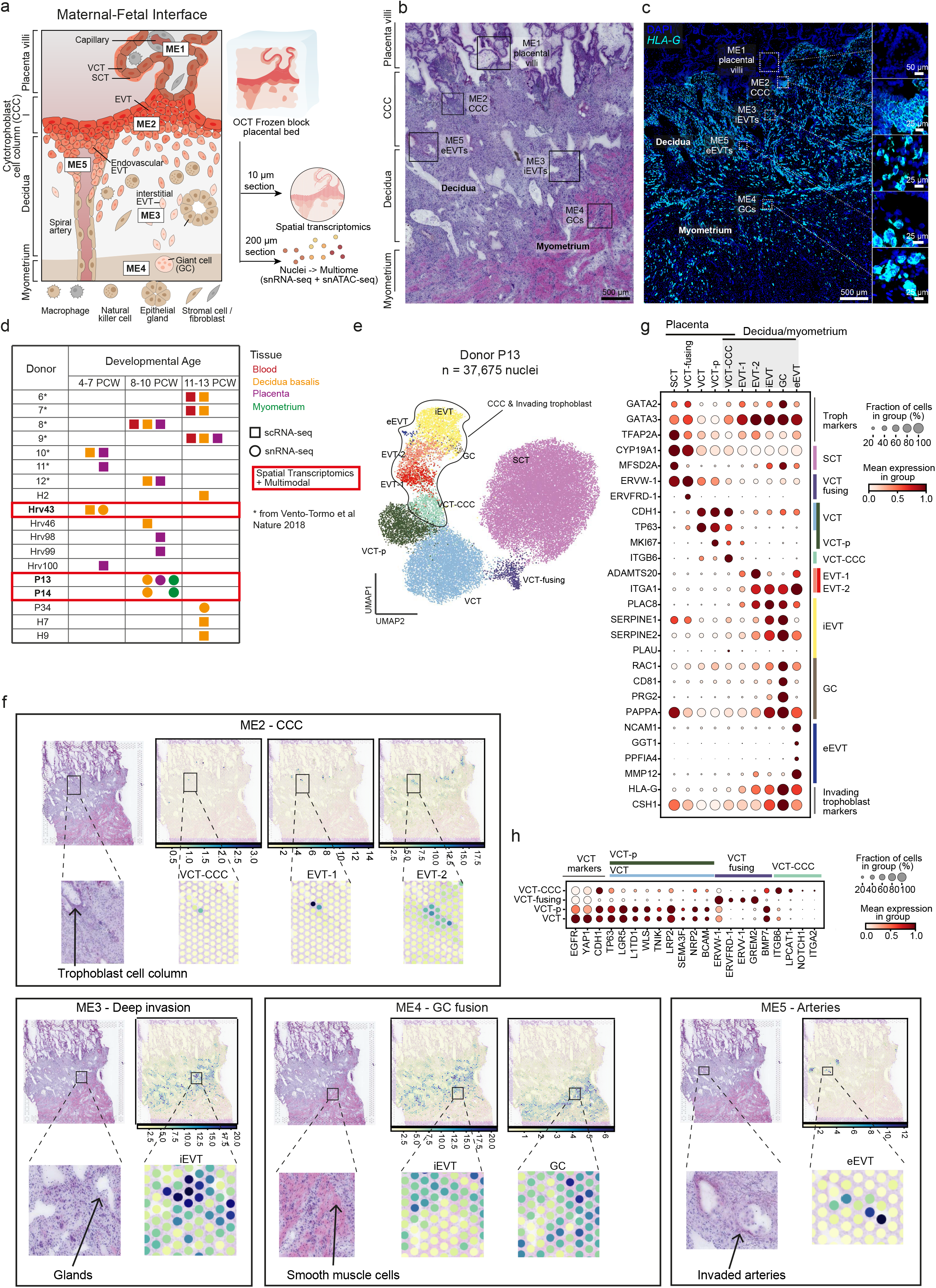
Trophoblast cell states in the early maternal-fetal interface. **a:** Schematic representation of the maternal-fetal interface (MFI) in early human pregnancy in the first trimester (left) and overview of experimental design of the study (right). **b:** Histological overview (H&E staining) of the implantation site of donor P13 (~ 8-9 post-conceptional weeks, PCW); black squares indicate trophoblast microenvironments in space. **c:** High-resolution imaging of a section of the placenta-decidua interface stained by in situ hybridization (smFISH) for HLA-G, illustrating the depth of invasion of EVTs into the uterus. Magnified insets (dashed squares) highlight the HLA-G-negative placental villi, and HLA-G-positive EVTs emerging from the CCC to invade the decidua and myometrium. **d:** Cohort composition split by gestational age (PCW) window representing tissues sampled from each donor and performed assays. Highlighted in red rectangles are the three donors whose tissues have been additionally profiled with spatial (Visium) and multiome assays. **e:** UMAP (uniform manifold approximation and projection) scatterplot of snRNA-seq of donor P13 trophoblast cell states in the maternal-fetal interface (n = 37,675 nuclei) coloured by cell state. **f:** Overview of spatial locations of invading trophoblast cell states in Visium spatial transcriptomics data of representative section of donor P13 (position of capture area is indicated with an arrow in Supplementary Fig. 1a). Cell type densities represented are derived with cell2location. Colorbars indicate the cell densities in a Visium spot. Invading trophoblast cell states are grouped based on the spatial microenvironment they represent. **g:** Dot plots show the variance-scaled, log-transformed expression of genes (X-axis) characteristic of trophoblast cell states (Y-axis) in donor P13 snRNA-seq data. **h:** Dot plot showing the variance-scaled, log-transformed expression of genes (X-axis) characteristic of villous cytotrophoblast (VCT) (Y-axis) in donor P13 snRNA-seq data. Syncytiotrophoblast (SCT), villous cytotrophoblast (VCT), cytotrophoblast cell column (CCC), proliferative (p), extravillous trophoblast (EVT), giant cells (GC), endovascular EVT (eEVT), single-cell RNA sequencing (scRNA-seq), single-nuclei RNA sequencing (snRNA-seq), microenvironment (ME).

We examined trophoblast heterogeneity in two steps. Firstly, we analysed the full-thickness implantation site from P13 (~9 PCW), as it contains both fetal (placenta) and maternal (decidua and myometrium) tissues on the same slide, and the tissue block is perfectly preserved and oriented **(Fig. 1e, Supplementary Fig. 3a)**. Secondly, we validated the trophoblast populations and their markers in the integrated dataset (~8-13 PCW) **(Supplementary Fig. 3b-c)**. Trophoblast subsets were annotated by considering canonical markers and their spatial location **(Fig. 1f-g, Supplementary Fig. 1a-b, 3c).** To assign spatial coordinates we used cell2location^24^, our probabilistic method to deconvolve the spatial voxels using our pre-defined snRNA-seq data **(Fig. 1f, Supplementary Fig. 1a-b)**. We then placed the trophoblasts into five pre-defined microenvironments (ME) in the tissue based on manual histological annotation.

In the placental villi (ME1), villous cytotrophoblasts (VCTs) fuse to form the overlying SCT layer that is in contact with maternal blood in the intervillous space. VCT subsets express high levels of the TFs *TP63* and *CDH1* in P13 donor **(Fig. 1g)** and the rest of the donors **(Supplementary Fig. 3d)**. VCT and VCT proliferative (VCT-p) upregulate known stem cell/progenitor markers (*LGR5, L1TD1, TP63*), WNT-signalling molecules (*WLS, TNIK, LRP2*), the *SEMA3F-NRP2* signalling complex, and the VCT marker *BCAM^25^* **(Fig. 1h, Supplementary Fig. 3e)**. We define an additional population of VCTs in the placental villi that we name VCT-fusing which the connectivity network PAGA^26^ indicates is an intermediate cell state between VCT and SCT (**Supplementary Fig. 3f)**. As VCT commit into VCT-fusing, they downregulate WNT (*WLS, TNIK, LRP2*) and BMP signals (*BMP7*, and upregulation of BMP antagonist *GREM2*), and upregulate the endogenous retroviral genes (*ERVW-1, ERVFRD-1, ERVV-1*) known to mediate trophoblast fusion **(Fig. 1h, Supplementary Fig. 3e)**^27^. Our nuclei isolation strategy allows capture of mature multinucleated SCT (*CYP19A1, MFSD2A*) not found in previous scRNA-seq studies^14,15^ **(Fig. 1g, Supplementary Fig. 3d)**.

Foci of cytotrophoblast cell columns arise from the VCTs that break through the SCT. These expand and form a shell around the conceptus which becomes discontinuous in the following weeks. EVTs begin to differentiate in cell columns but invasive EVTs only emerge when the villi attach to the maternal decidua as anchoring villi. In the trophoblast shell (ME2), we define an additional population of cytotrophoblast cell column VCT (VCT-CCC) **(Fig. 1f, Supplementary Fig. 1b).** VCT-CCC are proliferative and PAGA analysis shows they are likely to emerge from VCT/VCTp and give rise to EVT **(Supplementary Fig. 3f)**. This analysis suggests VCT is a common progenitor for both VCT-fusing, giving rise to SCT in the placenta, and VCT-CCC where EVTs emerge. As they commit into VCT-CCC, they downregulate WNT (*WLS, TNIK, LRP2*), upregulate *NOTCH1^28,29^*, perform an integrin shift (upregulating *ITGB6 and ITGA2*), and upregulate markers characteristic of the epithelial-mesenchymal transition (EMT) programme (*LPCAT1^30^*) **(Fig. 1h, Supplementary Fig. 3e)**. Expression of *NOTCH1* and *ITGA2* is characteristic of trophoblast progenitor cells located in the column niche^28,29^. In agreement with this finding, in ME2, VCT-CCC co-localise with EVT **(Fig. 1f, Supplementary 1b).** Altogether, our single-cell transcriptomics atlas defines the markers of a VCT population that can differentiate into VCT-fusing (progenitors of SCT) and is also likely to give rise to VCT-CCC (progenitors of EVTs).

## StOrder defines the invasion trajectory of EVTs into the decidua

To further investigate the EVT differentiation pathway as it arises from the cytotrophoblast cell columns of the anchoring villi to infiltrate maternal tissue, we leveraged both single-cell and spatial transcriptomics data using a three-step statistical framework, which we named StOrder **(see Methods)**. Firstly, StOrder builds a gene expression-based connectivity matrix (generated in our case by PAGA^26^) to establish putative connections between clusters. The values in this matrix are interpreted as pairwise similarity scores for cell states in the gene expression space **(Fig. 2a, Supplementary Fig. 4a)**. Secondly, StOrder generates a spatial covariance matrix that reflects pairwise proximity of trophoblast states that co-exist in space. To do so, StOrder takes as an input the estimated cell densities per spot (derived in our case with cell2location^24^) in Visium spatial transcriptomics data, and fits a Gaussian Process model that derives pairwise spatial covariance scores for all the cell state pairs **(Fig. 2a)**. This allows inference of which cell states are proximal in physical space and are likely gradually differentiating as they migrate. Third, StOrder reconstructs connections between cell states by summing the connectivity matrix (step 1) from single-cell transcriptomics data and the spatial covariance matrix (step 2) from the spatial data in a weighted manner **(Fig. 2a, Supplementary Fig. 4b-d)**. In sum, StOrder reconstructs the likely cell transitions in space by taking into account both the single-cell transcriptomics and the mini-bulk spatial transcriptomics data.

**Figure 2.**
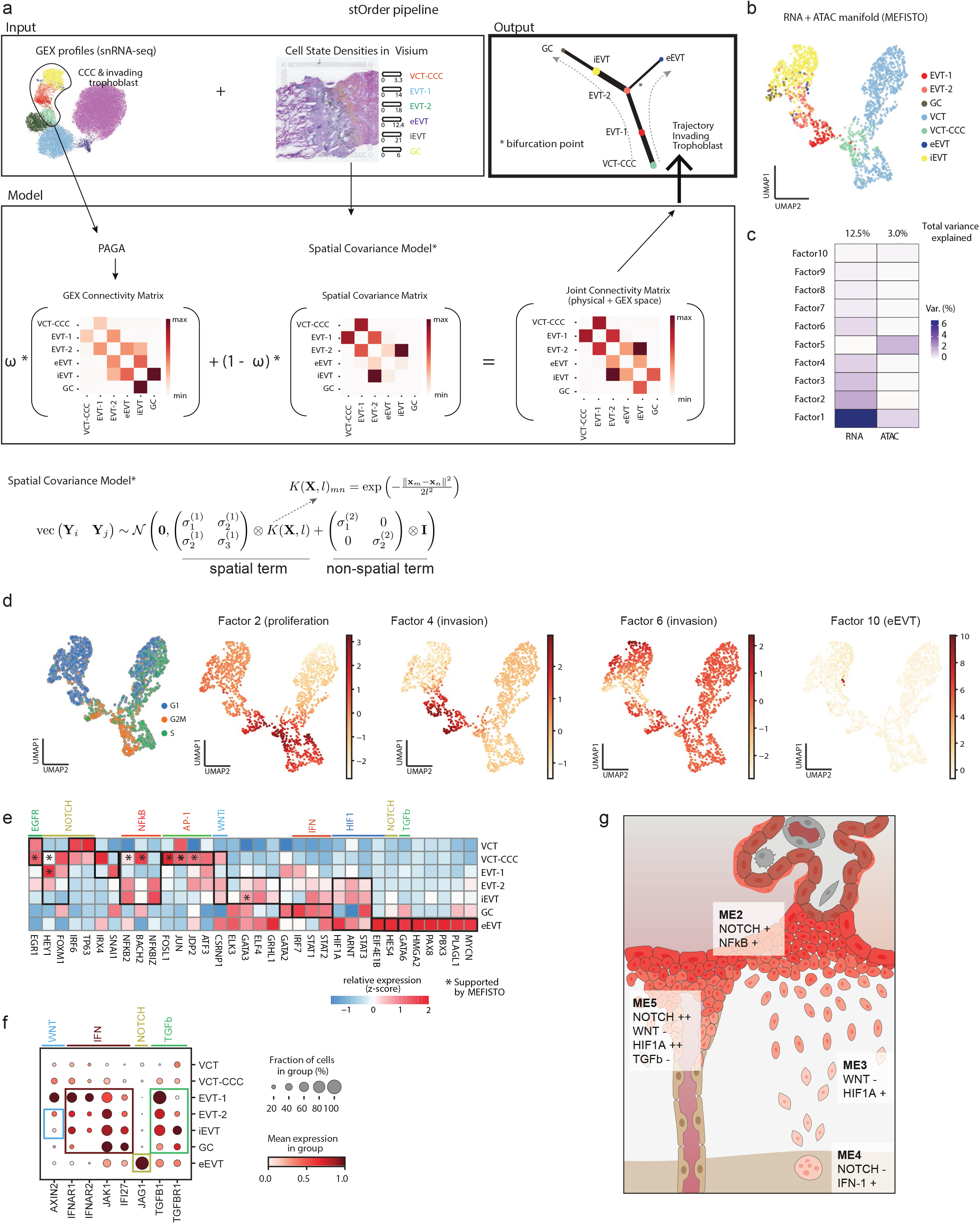
Regulatory programmes mediating extravillous trophoblast invasion. **a:** Schematic overview of StOrder approach representing the workflow of joint cell differentiation trajectory inference from gene expression and spatial data showing a representative tree with gene expression contribution of ω = 0.4. **b:** UMAP (uniform manifold approximation and projection) scatterplot of multiome (snRNA-ATACseq) data of invading trophoblast from donor P13 (n = 829) coloured by cell state. The manifold is calculated based on dimensionality reduction performed by MEFISTO (based on n=9 factors). **c:** Percentage of variance explained by each MEFISTO factor in each data modality. **d:** UMAP scatterplots of multiome (snRNA-ATACseq) data of invading trophoblast from donor P13 (n = 1605) as in **b** coloured by cell cycle phase and MEFISTO factor values for important selected factors **e:** Heatmap showing z-score of normalised, log-transformed and scaled expression of transcription factors (TF) relevant for trophoblast invasion in all donors. Y-axis indicates cell state, X-axis lists TFs. Differential expression is tested along invading trophoblast trajectory (as shown in Fig.2a) in a retrograde manner. Annotations of TFs on top of the heatmap are encoded as follows: asterix (*) = supported by MEFISTO; “a”= active TF. **f:** Dot plot showing the variance-scaled, log-transformed expression of genes (X-axis) of signalling molecules upregulated in EVT (Y-axis) in all donors. **g:** Schematic representation of signalling pathways in distinct microenvironments (see Figure 1a). Villous cytotrophoblast (VCT), cytotrophoblast cell column (CCC), proliferative (p), extravillous trophoblast (EVT), giant cells (GC), endovascular EVT (eEVT), microenvironment (ME), gene expression (GEX).

StOrder allowed us to resolve the most likely trajectory for the emergence and differentiation of invasive EVTs **(Fig. 2a)**. VCT-CCC are the precursors of EVTs-1 and EVTs-2, which co-localise with VCT-CCC in ME2 **(Fig. 1f, Supplementary 1b)**. EVT-1 are proliferative and closely related to VCT-CCC, while EVT-2 do not proliferate and have an early invasive phenotype, upregulating the metalloprotease *ADAMTS20* and the fibronectin-binding integrin *ITGA1* (**Fig. 1g, Supplementary Fig. 3d)**. EVT-2, located at the distal end of the columns of the anchoring villi, is identified as the bifurcation point **(Fig. 2a)**. EVT-2 can either transition into iEVTs, located in the invasion front of the decidua, or eEVTs, located inside the arteries. In agreement with eEVTs emerging from the tips of the columns, we detect spontaneous appearance of *NCAM1+* cells on a small number of EVTs in the cell columns **(Supplementary Fig. 5)**.

Highly invasive interstitial EVTs (iEVTs) are found in ME3, between decidual stromal and immune cells **(Fig. 1f)**. iEVTs upregulate *PLAC8^31^* and plasminogen activator inhibitors, *SERPINE1* and *SERPINE2*, with concomitant downregulation of plasminogen activator (*PLAU*) (**Fig. 1g, Supplementary Fig. 3d)**. iEVTs eventually fuse to form placental bed GCs deeper in the decidua and myometrium (ME4). GCs upregulate *RAC1* and *CD81*, both involved in myoblast fusion^32,33^, and the *PRG2-PAPPA* complex^34^ **(Fig. 1f-g, Supplementary Fig. 3c, Supplementary Fig. 1b)**. eEVTs, likely emerging from EVT-2, are present inside spiral arteries (ME5) **(Supplementary Fig. 3a)**. eEVTs express CD56 (*NCAM1*)^35,36^ and also upregulate the antioxidant enzyme *GGT1*, the liprin-associated member *PPFIA4*, and the metalloproteinase *MMP12* **(Fig. 1g, Supplementary Fig. 3d)**.

We next explored the regulatory programmes mediating EVT invasion by analysing the multimodal RNA-ATAC data **(Supplementary Fig. 4e-g)**. We applied our multifactorial method MEFISTO^37^ to donor P13 multimodal data, which contained the full spectra of VCT and EVT subsets **(Fig. 2b-c, Supplementary Fig. 4h-i)**. MEFISTO identified 10 latent factors that jointly explain 12.5% and 3% of the variance in the RNA expression data and the chromatin accessibility respectively **(Fig. 2c, Supplementary Fig. 4j, see Methods)**. Using a logistic regression approach, we define factors 2, 4, 6 and 10 as the main driving factors of the trophoblast trajectory **(Fig. 2d, Supplementary Fig. 4k-l)**. Factors 2, 4 and 6 explain changes along the main trophoblast invasion streak (VCTs-CCC through to GCs) (**Supplementary table 4**). Genes contributing strongly to these factors are *MKI67, CENPK* (cell cycle, factor 2); *CSF1R^38^, ADAM8, LAIR2^39^* (early trophoblast invasion, factor 4); *CALD1, COL21A1* (late trophoblast invasion, factor 6). Factor 10 captured eEVTs; the main genes contributing to this factor include *NCAM1, JAG1, ADORA1, EPHA1* and *HES4*.

## Transcription factor changes driving trophoblast fate during invasion

To identify the major regulatory programmes driving EVT differentiation, we extracted the transcription factors (TFs) that are differentially expressed and active along the EVT differentiation trajectory **(Supplementary Table 5)**. In addition, we included TFs whose binding motifs were enriched in top ATAC features of factors 2, 4, 6 and 10 in our multimodal analysis using MEFISTO **(Supplementary Table 5)**. As expected, activation of NOTCH (*HEY1, FOXM1, NOTCH1*) triggers differentiation of VCTs into VCT-CCC^28^ **(Fig. 2e)**. As previously shown, upregulation of *NOTCH1* may lead to the reduction of *IRF6* and *TP63* expression characteristic of VCT-CCC^28,40^. VCT-CCC upregulate the non-canonical NF-kB pathway (*NFKB2, BACH2*) and AP-1 factors (*FOSL1, JUN, JDP2, ATF3*), that may trigger the EMT program (e.g. upregulation *SNAI1*) **(Fig. 2e)**. Activation of the non-canonical NF-kB pathway is maintained throughout EVT differentiation, but there is upregulation of the NF-kB inhibitor (*NFkBIZ*) at the iEVT stage **(Fig. 2e)**. This could be a mechanism to avoid inflammation as EVTs invade^14^.

Decidual stromal cells secrete the WNT inhibitor DKK1^23^ and EVT invasion is marked by strong inhibition of WNT, with downregulation of the WNT target *AXIN2* and upregulation of the WNT repressor *CSRNP1* in iEVTs **(Fig. 2e-f)**. In addition, iEVTs upregulate TFs involved in cancer invasion (*ELK3-GATA3* complex^41^) and tumour suppressor genes (*ELF4, GRHL1*), in keeping with iEVTs being non-proliferative **(Fig. 2e)**. As iEVTs transition into GCs, they upregulate the type I IFN pathway, including TFs (*IRF7, STAT1, STAT2*), downstream transducers (*JAK1*),and targets (*IFI27*) **(Fig. 2e-f)**. These results suggest that type I IFN might play a role in GC fusion.

Following implantation and the formation of eEVT plugs in the spiral arteries, before 10 PCW the placenta is in a physiologically low-oxygen environment^42^. The hypoxia-inducible HIF1a pathway (*HIF1A, ARNT, STAT3*) is upregulated in both iEVTs and eEVTs but the HIF1A target *EIF4E1B^43^* is upregulated only in eEVTs, pointing to a role for this pathway in eEVT fate **(Fig. 2e-f)**. eEVTs also upregulate the NOTCH pathway (*HES4, JAG1*) and *GATA6*, both of them lowly expressed in iEVT. *GATA6* is known to affect vessels by suppressing autocrine TGFβ signalling^44^ and may have a similar role in this context, with both *TGFB1* and its receptor *TGFBR1* downregulated in eEVTs. This is different from iEVT, where both *TGFB1* and *TGFBR1* are upregulated. Additional TFs involved in the hypoxic environment in tumours and vessel transformation are upregulated in eEVTs, including *HMGA2^45^, PAX8^46^, PBX3^47^, PLAGL1^48^* and *MYCN*.

To summarise, our results point towards a key role for WNT inhibition, TGFβ and HIF1A activation in iEVT cell fate, while eEVT identity is marked by strong upregulation of NOTCH and HIF1A and strong downregulation of TGFβ signalling **(Fig. 2g)**.

## Invasive trophoblast subsets are recapitulated in tissue-derived placental organoids

We next explored if the cell-intrinsic regulatory programme that is triggered upon VCT-to-EVT differentiation is also present in our trophoblast organoids^11^. Our organoids are derived from primary placental cells and recapitulate the spontaneous fusion of VCT into SCT *in vitro*. Changing from trophoblast organoid medium (TOM) to EVT medium (EVTM) induces an invasive phenotype^11,49^. We differentiated organoids from six donors into EVTs and collected samples at 3h, 24h, 48h and 96h from the start of differentiation **(Fig. 3a-b)**. Organoids from both experiments were integrated into the same manifold and analysed in concert **(Fig.3c, Supplementary Fig. 6a)**. To define the identity of trophoblast states within the organoids, we first plotted the unique trophoblast markers identified in our *in vivo* atlas **(Supplementary Fig. 6b-c)**. Additionally, we projected the trophoblast *in vivo* reference data onto the *in vitro* trophoblast subsets by building a logistic regression classifier that we trained on the donor P13 trophoblast dataset^23^ **(Fig. 3d, Supplementary Fig. 6d)**.

**Figure 3.**
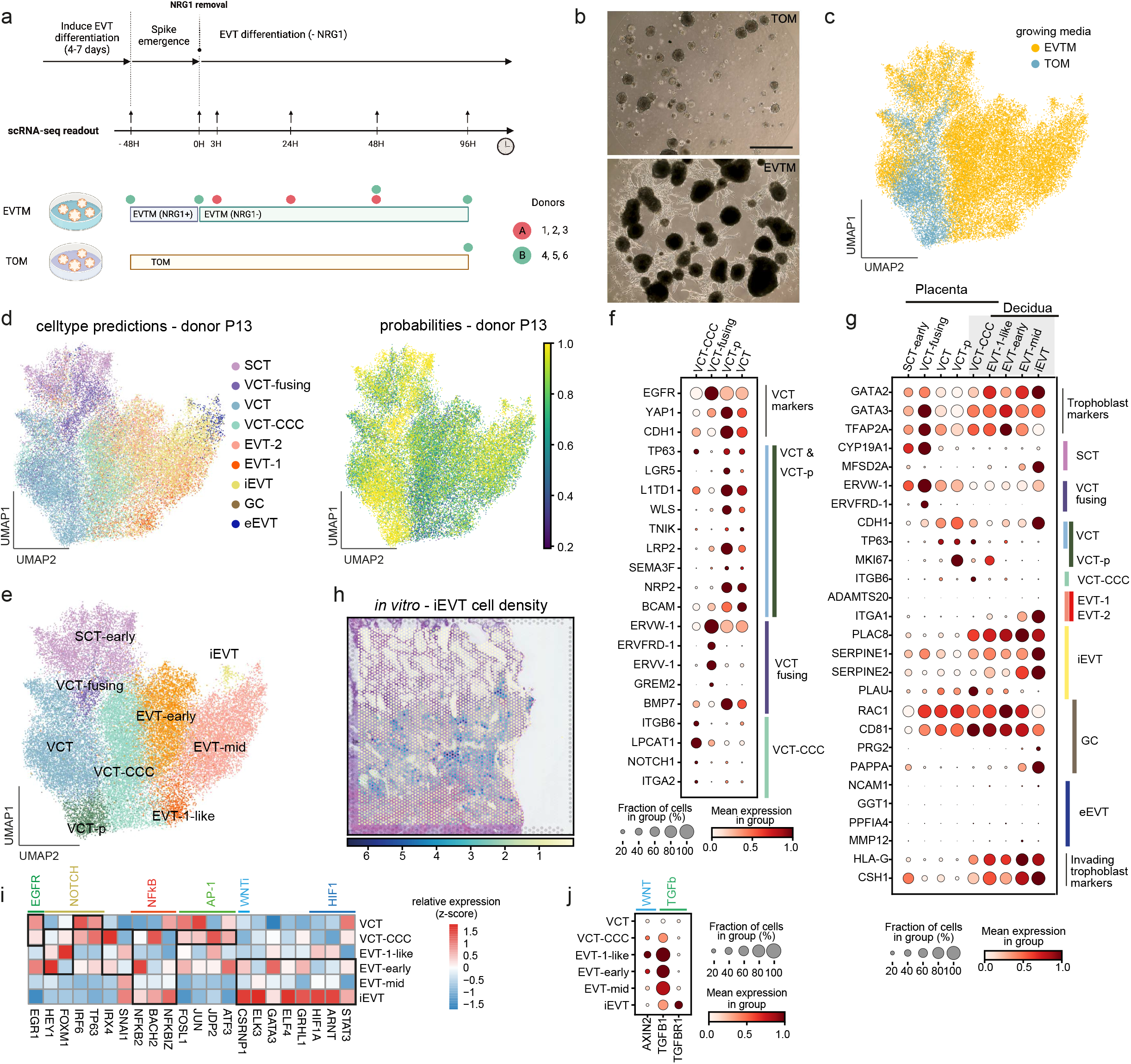
Regulatory programmes in primary derived trophoblast organoids. **a:** Schematic representation of the extravillous trophoblast differentiation experimental design, indicating time points and biological replicates (donors). **b:** Phase-contrast images of trophoblast organoids plated in a Matrigel droplet and exposed to TOM or EVTM. Scale bar is 1 mm. **c:** UMAP (uniform manifold approximation and projection) scatterplot coloured by growth medium. **d:** (Left) Predicted trophoblast subsets of placental organoids using a logistic classifier trained on P13 data. (Right) Logistic regression probabilities. **e:** UMAP scatterplot with final annotations of trophoblast subsets. **f:** Dot plot showing the variance-scaled, log-transformed expression of genes (X-axis) characteristic of villous cytotrophoblast (VCT) (Y-axis) in placental organoids. **g:** Dot plots show the variance-scaled, log-transformed expression of genes (X-axis) characteristic of trophoblast cell states (Y-axis) in placental organoids. **h:** Spatial locations of iEVTs in Visium spatial transcriptomics data of representative Visium section of donor P13 (position of capture area is indicated with an arrow in Supplementary Fig. 1A). Cell type densities represented are derived with cell2location trained on single-cell transcriptomics data of trophoblast organoids. Colorbars indicate the cell densities in a Visium spot. **i:** Heatmap showing z-score of normalised, log-transformed and scaled expression of transcription factors (TF) relevant for trophoblast invasion. Y-axis indicates cell state, X-axis lists TFs. Differential expression is tested along invading trophoblast trajectory (as shown in Fig.2a) in a retrograde manner. **j:** Dot plot showing the variance-scaled, log-transformed expression of genes (X-axis) of signalling molecules upregulated in EVT (Y-axis) in trophoblast organoids. Syncytiotrophoblast (SCT), villous cytotrophoblast (VCT), cytotrophoblast cell column (CCC), proliferative (p), extravillous trophoblast (EVT), giant cells (GC), endovascular EVT (eEVT), single-cell RNA sequencing (scRNA-seq), single-nuclei RNA sequencing (snRNA-seq).

We resolved the four VCT subsets identified *in vivo* in our trophoblast organoids. In the presence of TOM, the organoids were enriched in VCT (*LGR5, L1TD1, TP63, WLS, TNIK, LRP2, SEMA3F, NRP2, BCAM*) and VCT-fusing (*ERVW-1, ERVFRD-1, ERVV-1, GREM2*) **(Fig. 3c-f, Supplementary Fig. 6e)**. In our organoid dataset, SCT are *CYP19A1-low* and *MEFSD2A*-low, in agreement with the failure to capture fully differentiated multinucleated SCT by scRNA-seq **(Fig. 3g)**. A population of VCT-CCC (*ITGB6, LPCAT1, NOTCH1, ITGA2*) appeared only in the presence of EVTM **(Fig. 3f)**. EVTs emerge from VCT-CCC, suggesting that both *in vivo* and *in vitro*, VCTs have the potential to differentiate into either SCT or EVT lineages. These results suggest that cell fate shifts of VCT subsets are modulated by the culture conditions in *in vitro* trophoblast.

EVT populations arising in the presence of EVTM media were assigned as EVT. Despite some differences between the EVT subsets *in vivo* and *in vitro* (probability < 0.6), we find a small population in the organoids that corresponds to *in vivo*-iEVTs with a high probability score (probability > 0.8) **(Fig. 3d-e, Supplementary Fig. 6d)**. *In vitro*-iEVTs are enriched in later stages (48h and 96h), as expected, and are only present in two of the donors **(Supplementary Fig. 6e)**. Like their *in vivo* counterparts, iEVTs upregulate the plasminogen activator inhibitors *SERPINE1* and *SERPINE2* and downregulate *PLAU*. No expression of *NCAM1* is seen in differentiated organoid cultures, indicating the absence of eEVTs **(Fig. 3g)**. To further demonstrate the similarities between iEVTs *in vivo* and *in vitro*, we mapped *in vitro*-iEVTs onto the *in vivo* spatial transcriptomics data using cell2location^24^. *In vitro*-iEVTs exhibit a strong degree of localization to ME3 *in vivo* (Spearman rank-order correlation coefficient 0.91, p-value < 10e-308, two-sided test) (**Fig. 3h, Supplementary Fig. 6f-g, Supplementary Table 6**). This demonstrates the presence of invading iEVTs in our trophoblast organoid model and their suitability to study mechanisms modulating trophoblast invasion.

Finally, we used the organoids to define the intrinsic regulatory pathways mediating trophoblast invasion **(Fig. 3i-j)**. As in their *in vivo* counterparts, NOTCH-activated TFs (*HEY1, FOXM1, IRF6low*) and NF-kB TFs (*NFKB2, BACH2, JDP2, ATF3*) are present in VCT-CCC. The appearance of EVTs with an invasive phenotype is accompanied by downregulation of the WNT pathway (*CSRNP1, AXIN2* low), and upregulation of TFs involved in invasion (*ELK3-GATA3* complex^41^), tumour suppressor genes (*ELF4, GRHL1*) and the hypoxia inducible HIF1a pathway (*HIF1A, ARNT, STAT3*). Overall, we find the major programmes of EVT differentiation are conserved *in vivo* and *in vitro*. The subtle transcriptomic differences we encounter *in vitro* are likely to relate to the absence of maternal tissues. In addition, the lack of eEVTs in our culture indicates maternal factors absent in our cultures are required to define their identity.

## Modulation of trophoblast invasion by maternal cells in the decidua and myometrium

We next integrated single-cell and single-nuclei transcriptomics data from 18 donors to study how decidual maternal cells affect trophoblast invasion **(Fig. 1d, Fig. 4a, Supplementary Fig. 2e)**. We leveraged our tool CellPhoneDB v4^18^ to determine the ligand-receptor interactions that are enriched in the four decidual ME **(Fig. 1a, see Methods)**. We first focused on interactions mediating trophoblast invasion **(Fig. 4b)**. As previously described^14^, decidual natural killer cells (dNKs) interact with EVTs through multiple ligand-receptor pairs (*TGFBR1/2-TGFB, PVR-TIGIT, PVR-CD96, CCR1-CCL5, CSF1R-CSF1*). We find that the majority of these receptors are upregulated in EVT-2, close to the cytotrophoblast cell columns **(Fig. 4c)**. In this location *CSF1-CSF1R* interaction is enriched as shown by high-resolution multiplexed single molecule fluorescent in situ hybridisation (smFISH) **(Fig. 4d)**. *CSF1* is characteristic of dNK1^14,50^ and has a role in inducing tumour invasion^51^. Decidual macrophages (dM1 and dM2) are also likely to affect trophoblast invasion through expression of the chemokine *CXCL16*, which is known to interact with *CXCR6* upregulated in EVT-2 **(Fig. 4c)**. In addition, EVTs-2 express high levels of the guidance receptor *PLXND1* while its cognate ligand, *SEMA4C*, is characteristic of dM1.

**Figure 4.**
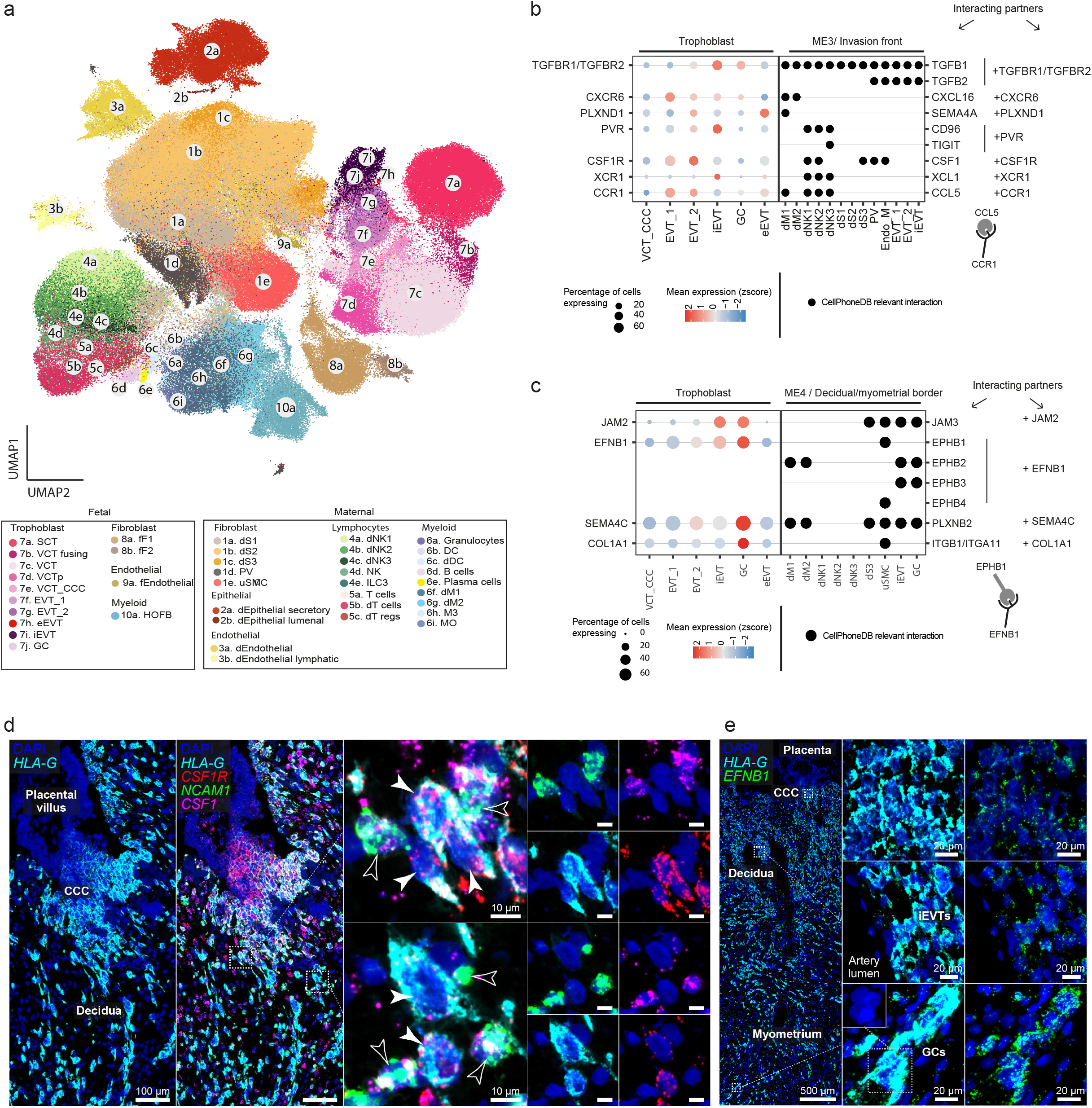
Cell-cell communication mediating extravillous trophoblast invasion. **a:** UMAP (uniform manifold approximation and projection) scatterplot of scRNA-seq and snRNA-seq of all donors described in Fig. 1d in the maternal-fetal interface (n = 350 815 cells and nuclei) coloured by cell state **b:** (Left) Dot plot showing the variance-scaled, log-transformed expression of genes (X-axis) of selected receptors upregulated in VCT-CCC (Y-axis) in trophoblast from all donors. (Right) Dot plot showing the presence (X-axis) of selected ligands in cells present in ME 3 (invasion front). Differential expression is tested along invading trophoblast trajectory (as shown in Fig. 2a) in a retrograde manner. **c:** (Left) Dot plot showing the variance-scaled, log-transformed expression of genes (X-axis) of selected receptors upregulated in EVT-1 and/or EVT-2 and or iEVT (Y-axis) in trophoblast from all donors. (Right) Dot plot showing the presence (X-axis) of selected ligands in cells present in ME 4 (decidual/myometrial border). Differential expression is tested along invading trophoblast trajectory (as shown in Fig. 2a) in a retrograde manner. **d:** (Left) High-resolution imaging of a section of the placenta-decidua interface stained by smFISH for *HLA-G*, highlighting EVTs invading the decidua from the CCC. (Centre) multiplexed co-staining with *NCAM1* (dNK marker), *CSF1* and cognate receptor *CSF1R*; dashed squares indicate areas shown magnified to right. (Right) solid and outlined arrows indicate neighbouring *CSF1R*-expressing EVTs and *CSF1*-expressing dNK cells, respectively. Representative image of samples from three donors. **e:** High-resolution imaging of a section of the placenta-decidua interface stained by multiplexed smFISH for *HLA-G* and *EFNB1*, demonstrating that expression of *EFNB1* is present throughout EVTs, including iEVTs, and elevated in GCs. Small inset at bottom-centre illustrates the multinucleated nature of GCs. Representative image of samples from two donors. Villous cytotrophoblast (VCT), cytotrophoblast cell column (CCC), extravillous trophoblast (EVT), giant cells (GC), endovascular EVT (eEVT).

The iEVTs invade as far as the inner third of the myometrium when they have fused into placental bed GCs^52^. GCs are probably no longer invasive because the receptors *CSF1R* and *PLXND1* are downregulated **(Fig. 4b).** In contrast, GCs upregulate adhesion molecules (*JAM2, EFNB1, SEMA4C*) whose cognate receptors are expressed by other iEVTs (*JAM3, EPHB2, EPHB3, PLXNB2*) that could be involved in cellular adhesion prior to their fusion^7^ **(Fig. 4c)**. Uterine smooth muscle cells (uSMCs) in the myometrium uniquely express *EPHB1* and *EPHB4* which bind to *EFNB1* upregulated in the iEVTs and GCs, possibly explaining their tropism towards the myometrium. We validated the expression of *EFNB1* in the GCs by multiplexed smFISH **(Fig. 4e)**. Altogether, we show a group of ligand/receptor pairs by which immune cells in the decidua may control invasion of EVTs and how these are downregulated in GCs.

## Decidual-trophoblast interactions mediating arterial remodelling

Trophoblast arterial transformation during early pregnancy is crucial for pregnancy success. Initially, there is medial destruction by iEVTs with replacement by acellular fibrinoid material^2,36,52^. Subsequently, eEVTs form a plug in the artery and partially replace the endothelium. This leads to loss of elasticity and dilation of the arteries essential to reduce the resistance to blood flow^52,53^. Making use of CellPhoneDB v4^18^, spatial transcriptomics and high-resolution microscopy, we next investigated how iEVTs and eEVTs jointly coordinate this process.

We mapped the interactions between perivascular cells (PVs)^14^ and iEVT. Expression of *EFNB1* by iEVTs could induce their tropism towards the arteries as only PVs express the cognate receptor, *EPHB6* **(Fig. 5a, Fig. 4e)**. We also find iEVTs upregulate specific cell signalling molecules (*PTPRS, NTN4*) whose cognate receptors are uniquely present in PVs (*NTRK3, NTRK2*) **(Fig. 5a)**. This family of neurotrophic tyrosine receptor kinases (NTRKs) has been associated with cellular survival in other contexts and these interactions are possibly involved in the appearance of ‘fibrinoid change’ in the arterial media due to death of PVs by iEVTs^2,36,52,53^. Using multiplexed smFISH, we validated the specific interaction between iEVTs (*HLA-G+*) expressing *PTPRS* and PVs (*MCAM+*) expressing *NTRK3* in the arteries **(Fig. 5b)**.

**Figure 5.**
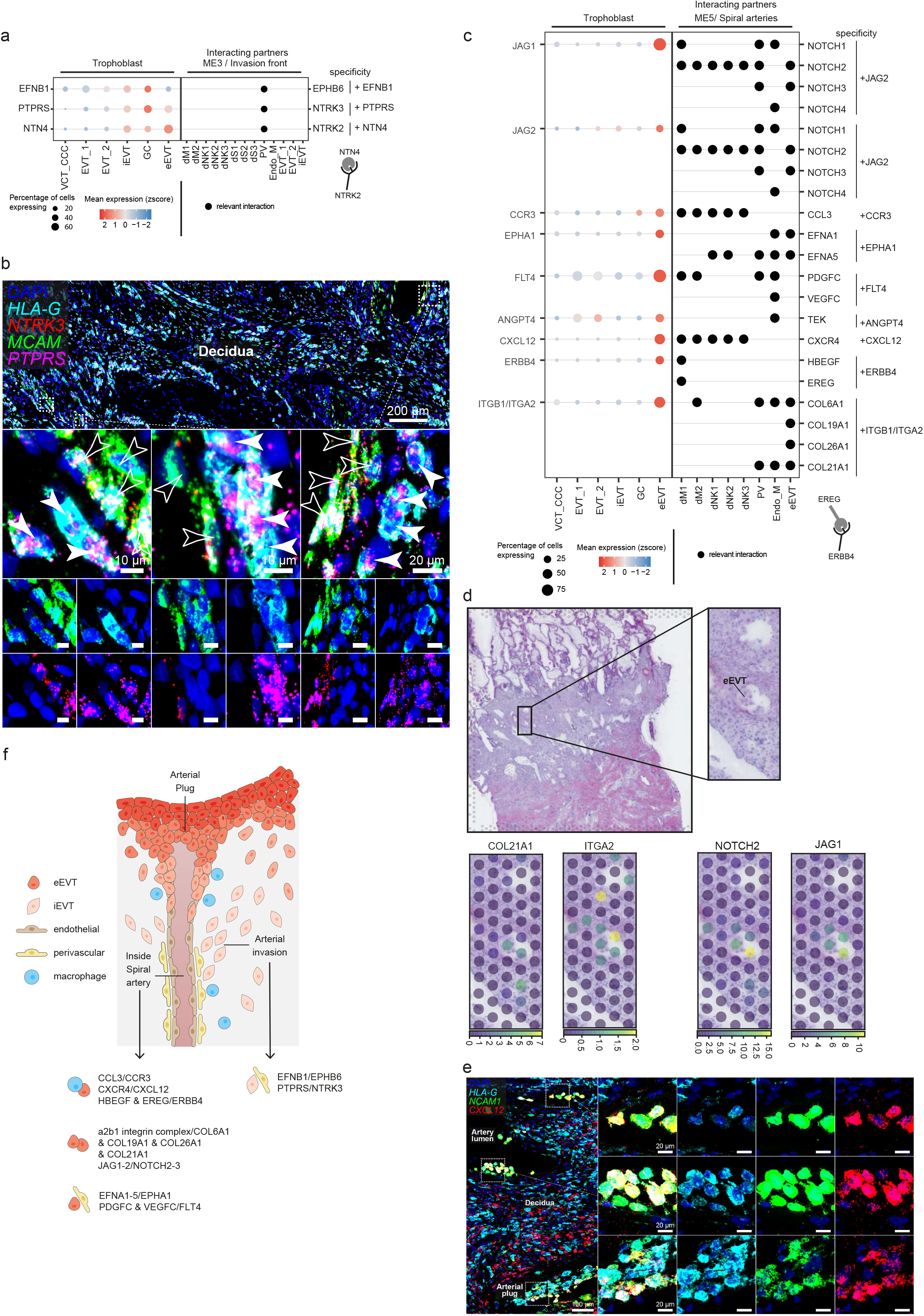
Cell-cell communication mediating arterial transformation during the first-trimester of pregnancy. **a:** (Left) Dot plot showing the variance-scaled, log-transformed expression of genes (X-axis) of selected receptors upregulated in iEVT(Y-axis) in trophoblast from all donors. (Right) Dot plot showing the presence (X-axis) of selected ligands in cells present in ME 3 (invasion front). Differential expression is tested along invading trophoblast trajectory (as shown in Fig. 2a) in a retrograde manner. **b:** (Top) High-resolution imaging of a section of decidua stained by multiplexed smFISH for *HLA-G, MCAM* (PV marker), *NTRK3* and its receptor *PTPRS;* dashed squares indicate areas shown magnified underneath. (Middle and below) solid and outlined arrows indicate neighbouring *PTPRS*-expressing EVTs and *NTRK3*-expressing dNK cells, respectively. Representative image of samples from three donors. **c:** (Left) Dot plot showing the variance-scaled, log-transformed expression of genes (X-axis) of selected receptors upregulated in VCT-CCC (Y-axis) in trophoblast from all donors. In the case of a complex, the expression corresponds to the least expressed subunit of the complex (ITGB1). (Right) Dot plot showing the presence (X-axis) of selected ligands in cells present in ME 5 (spiral arteries). Differential expression is tested along invading trophoblast trajectory (as shown in Fig. 2A) in a retrograde manner. **d:** Overview of spatial locations of invading trophoblast cell states in Visium spatial transcriptomics data of representative section of donor P13 (position of capture area is indicated with an arrow in Supplementary Fig. 1a). Cell type densities represented are derived with cell2location. Colorbars indicate the cell densities in a Visium spot. **e:** (Left) High-resolution imaging of a section of decidua stained by multiplexed smFISH for *HLA-G, NCAM1*, and *CXCL12*. Dashed squares highlight arteries containing *HLA-G+ NCAM1+* eEVTs expressing *CXCL12*, shown magnified to right. Representative image of samples from two donors. **f:** Schematic representation of the spiral arteries in early human pregnancy in the first trimester highlighting the novel interactions between PV-iEVT, endothelial-eEVT, immune-eEVT and eEVT-eEVT that we found in our dataset. Villous cytotrophoblast (VCT), cytotrophoblast cell column (CCC), extravillous trophoblast (EVT), giant cells (GC), endovascular EVT (eEVT).

The eEVT plugs limit maternal blood entering the intervillous space at high pressure before 8-10 PCW after which the full haemochorial circulation is established^54^. Our unbiased analyses of eEVTs allowed us to predict how the plugs are formed, revealing a specific ECM pattern that enables homotypic interactions. For example, *ITGB1* and *ITGA2* are expressed in eEVTs which together form the a2b1 complex that interacts with collagens specifically expressed by eEVTs (*COL6A1, COL19A1, COL26A1*, *COL21A1*) **(Fig. 5c)**. In addition, eEVTs upregulate both ligands (*JAG1* and *JAG2*) and receptors (*NOTCH2, NOTCH3*) that may stimulate active NOTCH signalling **(Fig. 5c)**. Using spatial transcriptomics, we visualised the presence of ECM components (e.g. *COL21A1-ITGA2*) and NOTCH interactions (e.g. *NOTCH2-JAG1*) in the arterial plug **(Fig. 5d)**.

Expression of chemokines and adhesion molecules could mediate interactions between eEVTs and the arterial wall. eEVTs upregulate *CCR3, EPHA1, CXCL12, ERBB4* whose ligands are expressed by endothelial (*EFNA1, EFNA5*) and immune (*CCL3, EFNA5, HBEGF, CXCR4, EREG*) cells. The unique expression of *CXCL12* by eEVTs was validated by smFISH **(Fig. 5e)**. eEVTs also upregulate *FLT4*, the receptor for *VEGFC*, upregulated by endothelial cells, and the growth factor *PDGFC* upregulated by endothelial cells, immune and PVs.

Altogether, by examining cell-cell interactions in the distinct trophoblast subsets, we map the cellular and molecular events mediating the transformation of the arteries during early pregnancy **(Fig. 5f)**.

## Discussion

In the post-implantation embryo, trophectoderm differentiates into trophoblast, the defining epithelial cells of the placenta that invade the uterus to transform the maternal arteries. Defective trophoblast invasion is the primary underlying cause of the great obstetric syndromes that include pre-eclampsia, fetal growth restriction, unexplained stillbirth, placental abruption and preterm labour^4^. We report new multiomics and spatial data, and develop a statistical framework (StOrder) that describes the complete trophoblast invasion trajectory during the first trimester of pregnancy. This includes the unbiased transcriptomics profile of eEVTs that move down inside the maternal arteries, and placental bed GCs, present deeper in the decidua and the inner myometrium.

We made use of a historical collection of pregnant hysterectomies at 8-10 PCW to delineate the landscape of the trophoblast at the implantation site, the place where fetal and maternal cells intermingle. The human implantation sites profiled in our study were collected more than 30 years ago and have been stored in liquid nitrogen. The discovery that these historical samples are so well preserved that we could use them for cutting edge single-cell transcriptomic analysis is important. Such samples are rare today owing to advances in clinical treatments that avoid hysterectomy during pregnancy. More broadly it shows how unanswered biological questions can be answered using such old samples. Our experimental design that combined consecutive sections for spatial transcriptomics and single-cell multiomics methods allowed us to integrate the molecular and cellular profiles with their spatial coordinates.

We define the transcriptomic profile of a VCT subset that can commit into SCT via a VCT-fusing intermediate, or invading EVTs via VCT located in the column niche (VCT-CCC). All these VCT subsets are present in our tissue-derived trophoblast organoid model, demonstrating that both i*n vivo* and *in vitro*, VCTs can give rise to both EVTs and SCT. This is in line with the observation that clonally derived trophoblast organoids can give rise to both cell types^55^. However, as VCT differentiate to EVT, there are subtle differences in the intermediate EVT subsets emerging *in vivo* and *in vitro*. Differences between *in vivo* and *in vitro* datasets could be explained by transient EVT populations not being captured in our *in vivo* dataset or because the influence of maternal serum and decidual tissues is lacking *in vitro*. Despite these small differences in the subsets emerging from the cytotrophoblast cell columns *in vivo* and *in vitro*, we do find an iEVT subset in the organoids that is present deeper in decidua *in vivo*. To compare this subset to *in vivo* iEVTs, we used the probabilistic method cell2location and can confirm that *in vitro*-iEVTs are equivalent to iEVTs present in ME3 from the *in vivo* spatial transcriptomics data. This means that our trophoblast organoid model can be used to explore questions such as how placental bed GCs form.

We developed StOrder, which uses both spatial and single-cell transcriptomics events to reconstruct the trajectory of trophoblast invasion and to identify bifurcation points. All code is available in our github and can be applied to other scenarios where lineage decisions are correlated with spatial changes. Our framework can be tailored to other spatial technologies by adapting specific parameters such as the distance between spots considered. StOrder pointed to EVT-2, located at the tips of the villi (or outer part of the cytotrophoblast shell), as the most likely precursors of eEVTs. Our result is consistent with previous histological observations suggesting that eEVTs arise from the cell columns or shell and move down the arterial lumen^35,36,56^. Using an approach that considers both TF expression and activity, we found eEVT identity is marked by a strong upregulation of NOTCH and HIF1A and a downregulation of TGFβ signalling. eEVTs are not found in the *in vitro* organoid model. This could be due to the absence of maternal endothelial cells or serum, or the non-hypoxic culture conditions.

Our systems biology approach has also allowed us to explore how the arterial transformation by both iEVTs and eEVTs is coordinated. Histological studies show that the medical destruction (fibrinoid change) is seen after iEVTs have encircled the arteries and it is only after this that eEVTs form a plug and then move down the artery in a retrograde manner to partially replace the endothelium^2,36,52^. We find selective interactions between iEVTs and PVs that may drive their tropism towards the arterial wall and mediate the destruction of arterial smooth muscle media. On the other hand, eEVTs have a specific ECM that could allow them to form the plug. There are also specific interactions with endothelial cells allowing adherence of eEVTs to them. These novel interactions add to our understanding of the communication between endothelial and eEVT cells^57^. The impact of defective arterial transformation in the later stages of pregnancy is well-described and underpins the great obstetric syndromes^6^. Our study increases understanding of these major pregnancy disorders that all have their origins in the first trimester^58^.

Deep trophoblast invasion into the uterus is exclusive to human and great apes^10^, and pre-eclampsia is only observed in humans^59^. Until now, our understanding of trophoblast invasion deep into the myometrium *in vivo* has mainly been limited to morphological and histological studies of archival specimens. Our study has identified markers of trophoblast invasion during a healthy pregnancy that can be compared to pathological conditions and cross referenced in genetic studies. Parallels are observed in other biological scenarios, such as cancer or tissue regeneration, and thus, some of the fundamental processes described in this work may extrapolate to other contexts. In addition, our bioinformatics tools will have a broader use for inferring spatial ordering of cells in other contexts, such as the tumour microenvironment. Finally, our roadmap of trophoblast differentiation can be used as a blueprint to design improved *in vitro* models that fully recapitulate the early stages after implantation.

## Supplementary Material

### Supplementary Figures

**Supplementary Fig. 1 Spatial transcriptomics of the implantation site.**

**a**: Histological overview (H&E staining) of donors P13, P14 and Hrv43 tissues with annotations of tissue regions. For the implantation site of donor P13 (~ 8-9 post-conceptional weeks, PCW, left); black squares indicate trophoblast microenvironments in space; faint grey squares (big) indicate positioning of tissue on Visium spatial transcriptomics capture areas; arrow indicates representative Visium section further explored in Fig. 1f.

**b**: Cell state locations (derived with cell2location) for representative Visium sections of donors P14 and Hrv43 highlighting relevant spatial trophoblast microenvironments.

Syncytiotrophoblast (SCT), villous cytotrophoblast (VCT), cytotrophoblast cell column (CCC), proliferative (p), extravillous trophoblast (EVT), giant cells (GC), endovascular EVT (eEVT), single-cell RNA sequencing (scRNA-seq), single-nuclei RNA sequencing (snRNA-seq), microenvironment (ME).

**Supplementary Fig. 2 Overview of analysis and quality control of coarse cell states in scRNA-seq and snRNA-seq data for the maternal-fetal interface.**

**a**: Overview of the computational pipeline implemented for analysis of scRNA-seq and snRNA-seq data.

**b-e**: (top) UMAP (uniform manifold approximation and projection) scatterplots of donors P13, P14, Hrv43 and all donors’ data (b-e respectively) for all recovered cell states, colored by coarse grain compartment annotation and important metadata labels: assay, sample (10X library), donor and developmental age. (bottom) Dot plots show the variance-scaled, log-transformed expression of genes characteristic of coarse grain compartment (X-axis) in donors profiled (Y-axis).

Maternal (m), fetal (f), natural killer (NK), innate lymphocytes (ILC)

**Supplementary Fig. 3 Overview of quality control of trophoblast cell states in scRNA-seq and snRNA-seq data for the maternal-fetal interface.**

**a**: UMAP (uniform manifold approximation and projection) scatterplots of donor P13 snRNA-seq data for all trophoblast cell states colored by assay, sample (10X library) and cell cycle phase of the nuclei.

**b:** UMAP scatterplot of integrated snRNA-seq and scRNA-seq of all donors’ trophoblast cell states in the maternal-fetal interface (n = 75,042 nuclei and cells) coloured by cell state

**c**: UMAP scatterplots of all donors’ scRNA-seq and snRNA-seq data for all trophoblast cell states colored by assay, sample (10X library), cell cycle phase of the cells/nuclei, donor and developmental age

**d**: Dot plots show the variance-scaled, log-transformed expression of genes (X-axis) characteristic of trophoblast cell states (Y-axis) in all donors.

**e**: Dot plots show the variance-scaled, log-transformed expression of genes (X-axis) characteristic of trophoblast cell states (Y-axis) in all donors.

**f:** Results of PAGA trajectory inference of all trophoblast cell states in donor P13 snRNA-seq data (left: main manifold, center: denoised PAGA manifold, right: PAGA reconstruction of putative trajectory tree for all trophoblast cell states). For the purpose of this analysis all EVTs have been united in annotation under ‘EVT’ label.

Syncytiotrophoblast (SCT), villous cytotrophoblast (VCT), cytotrophoblast cell column (CCC), proliferative (p), extravillous trophoblast (EVT), giant cells (GC), endovascular EVT (eEVT), single-cell RNA sequencing (scRNA-seq), single-nuclei RNA sequencing (snRNA-seq).

**Supplementary Fig.4 Multimodal analysis of extravillous trophoblast invasion.**

**a**: (Left) Main UMAP (uniform manifold approximation and projection) scatterplot and (right) denoised manifold used for PAGA trajectory inference of all trophoblast cell states in donor P13 snRNA-seq data.

**b:** PAGA reconstruction of putative trajectory tree for all extravillous trophoblast cell states. This corresponds to the trajectory inferred using ω= 1 in StOrder.

**c**: Reconstruction of putative invading trophoblast trajectory tree based on both gene expression and spatial data (range of ω ∈ [0.3, 0.48] in stOrder approach). stOrder was performed on donor P13 snRNA-seq and donors P13, P14 and Hrv43 spatial locations for invading trophoblast and VCT_CCC cell states.

**d**: Reconstruction of putative invading trophoblast trajectory tree based solely on spatial data (ω=0 in stOrder approach). stOrder was performed on donor P13 snRNA-seq and donors P13, P14 and Hrv43 spatial locations for invading trophoblast and VCT_CCC cell states.

**e:** Overview of the computational pipeline implemented for analysis of multimodal data.

**f:** UMAP scatterplots of integrated multimodal data from donors P13, P14 and hrv43. Data annotated based on the snRNA-seq annotation.

**g:** UMAP scatterplots coloured by donor, sample and unbiased clustering

**h:** UMAP scatterplots of trophoblast cell states.

**i:** (Left) UMAP scatterplot of multiome (snRNA-ATACseq) data of invading trophoblast from donor P13 (n = 829) coloured by sample. The manifold is calculated based on dimensionality reduction performed by MEFISTO (model with n=9 factors). (Right) Scatterplot of UMAP coordinates obtained from the RNA expression data that were used as covariates for MEFISTO. Each dot corresponds to a cell coloured by lineage assignment.

**j:** Estimated smoothness along differentiation.

**k:** Learnt correlation structure for each latent factor.

**l**: Gene set (RNA, left) enrichment analysis overview of MEFISTO factor 2.

**m:** Peak set (ATAC, right) enrichment analysis overview of MEFISTO factor 10.

Villous cytotrophoblast (VCT), cytotrophoblast cell column (CCC), proliferative (p), extravillous trophoblast (EVT), giant cells (GC), endovascular EVT (eEVT), dendritic cells (DC), lymphatic (l), maternal (M), Hofbauer cells (HOFB), innate lymphocytes (ILC), macrophages (M), monocytes (MO), natural killer (NK), perivascular (PV), decidua (d), epithelial (epi), stromal (S), fibroblasts (F), uterine smooth muscle cells (uSMC).

**Supplementary Fig. 5. *NCAM1+* eEVTs emerging from the cytotrophoblast cell column**

(Top) High-resolution imaging of sections of the placenta-decidua interface stained by multiplexed smFISH for *HLA-G* and *NCAM1*. (Middle) magnified insets highlight cytotrophoblast cell columns and solid arrows indicate *HLA-G+ NCAM1+* cells (nascent eEVTs) shown magnified below (bottom). Images of samples from two donors shown.

**Supplementary Fig. 6. Benchmark of primary-derived placental organoids.**

**a:** UMAP (uniform manifold approximation and projection) scatterplots of 6 organoid donors colored by donor, time-point and cell cycle.

**b:** UMAP scatterplot coloured by unbiased clustering using louvain.

**c:** Dot plot showing the variance-scaled, log-transformed expression of genes (X-axis) of main trophoblast subsets (Y-axis) on each of the cells identified by unbiased clustering (B).

**d:** Bar plot showing the proportion of predicted cell states by our logistic regression model on each of the identified clusters (B).

**e:** Bar plot showing the proportion of final cell states identified on each donor (left) and time-point (right).

**f:** Overview of spatial locations of EVT-mid and iEVT subsets in 10X Visium spatial transcriptomics data in Visium sections of donor P13. Cell type densities represented are derived with cell2location with single-cell transcriptomics data from the organoids used as a reference. Colorbars indicate the cell state density in a Visium spot.

**g:** Scatterplot of cell densities derived by cell2location of *in vitro* iEVT (X-axis, using single-cell transcriptomics of trophoblast organoids) vs *in vivo* iEVT (Y-axis, using single-nucleus transcriptomics of donor P13) cell states in donor P13 Visium sections WS_PLA_S9101764, WS_PLA_S9101765, WS_PLA_S9101766 and WS_PLA_S9101767. In red is the trend line representing Spearman rank-order correlation (R = 0.91, p-value < 10e-308, two-sided test) between values of cell densities of *in vivo* iEVT and *in vitro* iEVT. Syncytiotrophoblast (SCT), villous cytotrophoblast (VCT), cytotrophoblast cell column (CCC), proliferative (p), extravillous trophoblast (EVT), interstitial EVT (iEVT), giant cells (GC), endovascular EVT (eEVT).

## Acknowledgements

This publication is part of the Human Cell Atlas. We gratefully acknowledge the Sanger Cellular Generation and Phenotyping (CGaP) Core Facility and the Sanger Core Sequencing pipeline for support with sample processing and sequencing library preparation. Ricard Argelaguet for his help on MOFA/MEFISTO analysis. Vitalii Kleshchevnikovfor this help on cell2location. Stijn van Dongen, Martin Prete and Simon Murray for their help on running pipelines and web portal support. Tarryn Porter for her help on preparing the libraries for spatial transcriptomics. Antonio Garcia for graphical images. Aidan Maartens for proofreading. Placental material was provided by the Joint MRC-Human Cell Atlas (MR/S036350/1). The authors are grateful to patients for donating tissue for research. We thank D. Moore and M. Maquinana and staff at Addenbrooke’s Hospital, Cambridge, UK.

## Contributions

A.M collected and analysed the histology of the samples; E.P and R.H performed the nuclei experiments; K.R and E.T performed the spatial transcriptomics analyses with help of C.I.M and I.K; M.A.S derived all organoid lines, performed all organoid culturing and prepared all time-point collections; A.A, B.V, L.G-A and K-T analysed all the data; A.A and I.K developed StOrder; A.A, M.Y.T and R.V.T interpreted the data with contribution of A.M, M.A.S and K.R; R.V.T and O.S supervised the bioinformatics analyses; R.V.T and O.B supervised the *in vivo* and genomics work; M.Y.T supervised the *in vitro* work; R.V.T wrote the manuscript with contributions from K.R, A.M, M.A.S, and A.A; the final version of the manuscript has been approved by all the authors.

## Data availability

Datasets are being uploaded into EMBL-EBI ArrayExpress and can now be accessed at https://www.reproductivecellatlas.org/mfi/. All codes used for data analysis are available from https://github.com/ventolab/MFI

## Supplementary Tables

**Supplementary Table 1. Supplementary_Table_1.xlsx (separate file)**

**Metadata of samples. (A)** 10X scRNA-seq libraries from human donors. **(B)** 10X snRNA-seq libraries from human donors. **(C)** 10X cell-coupled snRNA/ATAC-seq (multiome) libraries from human donors. **(D)** 10X Visium spatial transcriptomics libraries from human donors. Sample id = 10x reaction; Donor = donor ID; Stage_PCW = post-conceptional weeks; TP = type of pregnancy termination (Med: medical; Sur: surgical or Hys: hysterectomy)

**Supplementary Table 2. Supplementary_Table_2.xlsx (separate file)**

**Quality control of samples for each 10X RNA library in our maternal-fetal interface atlas. (A)** Summary statistics from 10X Cell Ranger 3.0.2 for scRNA-seq samples. **(B)** Summary statistics from 10X Cell Ranger 3.0.2 for snRNA-seq samples. **(C)** Summary statistics from 10X Cell Ranger ARC 1.0.1 for multiome samples. **(D)** Summary statistics from 10X Space Ranger 1.1.0 Visium spatial transcriptomics samples.

**Supplementary Table 3. Supplementary_Table3.xlsx (separate file)**

**Annotation summary for each sample.** Number of cells/nuclei (droplets) per coarse cell state in scRNA-seq, snRNA-seq and multiome samples of donors P13, P14, Hrv43 and all donors dataset.

**Supplementary Table 4. Supplementary_Table4.xlsx (separate file)** Variance explained (R2 column) in the MEFISTO model by each factor in each modality (RNA or ATAC).

**Supplementary Table 5. Supplementary_Table5.xlsx (separate file)**

TF analysis along trophoblast trajectory. Table containing the multiple TF measurements in the *in vivo* analysis used to prioritise TF relevant for trophoblast differentiation of all TFs **(A)** and selected TFs **(B)**. All tests are performed by comparing the newly emerged cell type against the pseudo-ancestor. Columns across table indicate: TF = transcription factor; cluster = cell type; regulation_sign = whether up or downregulation is tested; Avg_expr = average log-transformed normalised expression within the cell type; is_DE_limma = ‘yes’ if it is a differentially expressed TF (FDR < 0.05; limma); is_DA_dorothea = ‘yes’ if it is a differentially activated TF (FDR < 0.05; Wilcoxon test); is_DA_chromVar = ‘yes” if the TF binding motifs are differentially accessible (FDR < 0.05; Wilcoxon test); is_DA_MEFISTO ‘yes” if the TF binding motifs are differentially accessible in the regions linked to MEFISTO factor (FDR < 0.05; Wilcoxon test); if_DA_MEFISTO_factor = MEFISTO factor associated; is_DE_and_DA = ‘yes’ if the TF is differentially expressed and differentially activated according to any other measure.

**Supplementary Table 6. Supplementary_Table6.xlsx (separate file)**

Cell2location cell density values of *in vitro* iEVTs (using single-cell transcriptomics of trophoblast organoids) and *in vivo* iEVTs (using single-nucleus transcriptomics of donor P13) cell states in donor P13 Visium sections WS_PLA_S9101764, WS_PLA_S9101765, WS_PLA_S9101766 and WS_PLA_S910176.

**Supplementary Table 7. Supplementary_Table7.xlsx (separate file)**

Trophoblast interactions enriched by microenvironment (ME) using CellPhoneDB. **(A)** ME2 = cytotrophoblast cell column. **(B)** ME3 = Invasion front. **(C)** ME4 = Decidual/myometrial border. **(D)** ME5 = Spiral arteries.

**Supplementary Table 8. Supplementary_Table8.xlsx (separate file)**

Probes used for multiplexed RNAscope smFISH.

## Materials and methods

### Patient samples

Tissue samples used for this study were obtained with written informed consent from all participants in accordance with the guidelines in The Declaration of Helsinki 2000.

Placental and decidual samples used for the *in vivo* and *in vitro* profiling were obtained from elective terminations from:

- The MRC and Wellcome-funded Human Developmental Biology Resource (HDBR, http://www.hdbr.org), with appropriate maternal written consent and approval from the Fulham Research Ethics Committee (REC reference 18/LO/0822) and Newcastle & North Tyneside 1 Research Ethics Committee (REC reference 18/NE/0290). The HDBR is regulated by the UK Human Tissue Authority (HTA; www.hta.gov.uk) and operates in accordance with the relevant HTA Codes of Practice.
- Addenbooke’s Hospital (Cambridge) under ethical approval from the Cambridge Local Research Ethics Committee (04/Q0108/23), which is incorporated into The overarching ethics permission given to the Centre for Trophoblast Research biobank for the “Biology of the Human Uterus in Pregnancy and Disease Tissue Bank” at the University of Cambridge under ethical approval from the East of England-Cambridge Central Research Ethics Committee (17/EE/0151) and from the London-Hampstead Research Ethics Committee (20/LO/0115).

Placental/decidual blocks (P13, P14 and P34) were collected prior to 1 September 2006 and consent for research use was not obtained. These samples are considered ‘Existing Holdings’ under the Human Tissue Act and as such were able to be used in this project.

All samples profiled were histologically normal.

### Tissue cryopreservation

Fresh tissue samples of human implantation sites were embedded in cold OCT medium and flash frozen using a dry ice-isopentane slurry. Protocol available at protocols.io^60^.

Quality of archival frozen tissue samples was assessed by extraction of RNA from cryosections using the QIAGEN RNeasy Mini Kit, according to the manufacturer’s instructions including on-column DNase I digestion. RNA quality was assayed using the Agilent RNA 6000 Nano Kit. All samples processed for Visium and single-nuclei had RIN values greater than 8.7.

### Single-nuclei extraction

Single-nuclei suspensions were isolated from frozen tissue sections when performing multiomic snRNA-seq/scATAC-seq and snRNA-seq, following manufacturer’s instructions. For each OCT-embedded sample, 400 μm of tissue was prepared as 50 μm cryosections, which were paused in a tube on dry ice until subsequent processing. Nuclei were released via Dounce homogenisation as described in detail at protocols.io^61^.

### Tissue processing

We used the previous protocol optimised for the decidual-placental interface^14^. In short, decidual tissues were enzymatically digested in 15 ml 0.4 mg/ml collagenase V (Sigma, C-9263) solution in RPMI 1640 medium (Thermo Fisher Scientific, 21875-034)/10% FCS (Biosfera, FB-1001) at 37 °C for 45 min. The supernatant was diluted with medium and passed through a 100-μm cell sieve (Corning, 431752) and then a 40-μm cell sieve (Corning, 431750). The flow-through was centrifuged and resuspended in 5 ml of red blood cell lysis buffer (Invitrogen, 00-4300) for 10 min. Placental villi were scraped from the chorionic membrane using a scalpel and the stripped membrane was discarded. The resultant villous tissue was enzymatically digested in 70 ml 0.2% trypsin 250 (Pan Biotech P10-025100P)/0.02% EDTA (Sigma E9884) in PBS with stirring at 37 °C for 9 min. The disaggregated cell suspension was diluted with medium and passed through a 100-μm cell sieve (Corning, 431752). The undigested gelatinous tissue remnant was retrieved from the gauze and further digested with 10–15 ml collagenase V at 1.0 mg/ml (Sigma C9263) in Ham’s F12 medium/10% FBS with gentle shaking at 37 °C for 10 min. The disaggregated cell suspension was diluted with medium and passed through a 100-μm cell sieve (Corning, 431752). Cells obtained from both enzyme digests were pooled together and passed through a 100-μm cell sieve (Corning, 431752) and washed in Ham’s F12. The flow-through was centrifuged and resuspended in 5 ml of red blood cell lysis buffer (Invitrogen, 00-4300) for 10 min.

### Trophoblast organoid cultures

In total, six trophoblast organoids were grown and differentiated into EVT as previously described^11,55^. To differentiate trophoblast organoids into EVT, organoids were cultured with trophoblast organoid media (TOM) for ~3-4 days and transferred into EVT media 1 (+NRG1) for ~4-7 days. Once trophoblasts initiate their commitment into EVT (spike emergence), EVT media 2 (-NRG1) is added for 4 days. Donors were differentiated and collected in batches of three that were multiplexed on the same 10x-genomics reaction. Samples for donors 1, 2 and 3 were collected at 3 hours (h), 24h and 48h after addition of EVTM media 2, while samples for donors 4, 5 and 6 were collected at 48h before, and then 0h, 48h and 96h after, addition of EVTM media 2. Organoids grown in trophoblast organoid media (TOM) media were also collected as a control at 96h.

Media composition was as described previously^11,55^:

TOM = Advanced DMEM/F12, N2 supplement (at manufacturer’s recommended concentration), B27 supplement minus vitamin A (at manufacturer’s recommended concentration), Primocin 100 μg/mL, *N*-Acetyl-L-cysteine 1.25 mM, L-glutamine 2 mM, recombinant human EGF 50 ng/mL, CHIR99021 1.5 μM, recombinant human R-spondin-1 80 ng/mL, recombinant human FGF-2 100 ng/mL, recombinant human HGF 50 ng/mL, A83-01 500 nM, prostaglandin E2 2.5 μM, Y-27632 5 μM.

EVT media 1 = Advanced DMEM/F12, L-glutamine 2 mM, 2-mercaptoethanol 0.1 mM, penicillin/streptomycin solution 0.5% (vol/vol), BSA 0.3% (vol/vol), ITS-X supplement 1% (vol/vol), NRG1 100 ng/mL, A83-01 7.5 μM, Knockout serum replacement 4% (vol/vol)

EVT media 2 = Advanced DMEM/F12, L-glutamine 2 mM, 2-mercaptoethanol 0.1 mM, penicillin/streptomycin solution 0.5% (vol/vol), BSA 0.3% (vol/vol), ITS-X supplement 1% (vol/vol), A83-01 7.5 μM, Knockout serum replacement 4% (vol/vol) ie: EVT medium cat. no. 1 without NRG1. Store the medium at 4°C for up to 1 week.

### Haematoxylin and eosin (H&E) staining and imaging

Fresh frozen sections were removed from −80°C storage and air dried before being fixed in 10% neutral buffered formalin for 5 minutes. After rinsing with deionised water, slides were stained in Mayer’s haematoxylin solution for 90 seconds. Slides were completely rinsed in 4-5 washes of deionised water, which also served to blue the haematoxylin. Aqueous eosin (1%) was manually applied onto sections with a pipette and rinsed with deionised water after 1-3 seconds. Slides were dehydrated through an ethanol series (70%, 70%, 100%, 100%) and cleared twice in 100% xylene. Slides were coverslipped and allowed to air dry before being imaged on a Hamamatsu Nanozoomer 2.0HT digital slide scanner.

### Multiplexed smFISH and high-resolution imaging

Large tissue section staining and fluorescent imaging was conducted largely as described previously^62^. Sections were cut from fresh frozen samples embedded in OCT at a thickness of 10-16 μm using a cryostat, placed onto SuperFrost Plus slides (VWR) and stored at −80°C until stained. Tissue sections were processed using a Leica BOND RX to automate staining with the RNAscope Multiplex Fluorescent Reagent Kit v2 Assay (Advanced Cell Diagnostics, Bio-Techne), according to the manufacturers’ instructions. Probes may be found in Supplementary Table 8. Prior to staining, fresh frozen sections were post-fixed in 4% paraformaldehyde in PBS for 6-8 hours, then dehydrated through a series of 50%, 70%, 100%, and 100% ethanol, for 5 minutes each. Following manual pre-treatment, automated processing included heat-induced epitope retrieval at 95°C for 15 minutes in buffer ER2 and digestion with Protease III for 15 minutes prior to probe hybridisation. Tyramide signal amplification with Opal 520, Opal 570, and Opal 650 (Akoya Biosciences) and TSA-biotin (TSA Plus Biotin Kit, Perkin Elmer) and streptavidin-conjugated Atto 425 (Sigma Aldrich) was used to develop RNAscope probe channels.

Stained sections were imaged with a Perkin Elmer Opera Phenix Plus High-Content Screening System, in confocal mode with 1 μm z-step size, using a 20X (NA 0.16, 0.299 μm/pixel) or 40X (NA 1.1, 0.149 μm/pixel) water-immersion objective. Channels: DAPI (excitation 375 nm, emission 435-480 nm), Atto 425 (ex. 425 nm, em. 463-501 nm), Opal 520 (ex. 488 nm, em. 500-550 nm), Opal 570 (ex. 561 nm, em. 570-630 nm), Opal 650 (ex. 640 nm, em. 650-760 nm).

#### Image stitching

Confocal image stacks were stitched as two-dimensional maximum intensity projections using proprietary Acapella scripts provided by Perkin Elmer.

### 10x Genomics Chromium GEX library preparation and sequencing

For the scRNA-seq experiments, cells were loaded according to the manufacturer’s protocol for the Chromium Single Cell 3’ Kit v3.0, v3.1 and 5’ v1.0 (10X Genomics). Library preparation was carried out according to the manufacturer’s protocol to attain between 2,000 and 10,000 cells per reaction. Libraries were sequenced, aiming at a minimum coverage of 20,000 raw reads per cell, on the Illumina HiSeq 4000 or Novaseq 6000 systems; using the sequencing format;

a. read 1: 26 cycles; i7 index: 8 cycles, i5 index: 0 cycles; read 2: 98 cycles
b. read 1: 28 cycles; i7 index: 8 cycles, i5 index: 0 cycles; read 2: 91 cycles
c. read 1: 28 cycles; i7 index: 10 cycles; i5 index: 10 cycles; read 2: 90 cycles (v3.1 dual)

For the multimodal snRNA-seq/scATAC-seq experiments, cells were loaded according to the manufacturer’s protocol for the Chromium Single Cell Multiome ATAC + Gene Expression v1.0 to attain between 2,000 and 10,000 cells per well. Library preparation was carried out according to the manufacturer’s protocol. Libraries for scATAC-seq were sequenced on Illumina NovaSeq 6000, aiming at a minimum coverage of 10,000 fragments per cell, with the following sequencing format; read 1: 50 cycles; i7 index: 8 cycles, i5 index: 16 cycles; read 2: 50 cycles.

### 10x Genomics Visium library preparation and sequencing

Ten micron cryosections were cut and placed on Visium slides, then processed according to the manufacturer’s instructions. Briefly, sections were fixed with cold methanol, stained with haematoxylin and eosin and imaged on a Hamamatsu NanoZoomer S60 before permeabilisation, reverse transcription and cDNA synthesis using a template-switching protocol. Second-strand cDNA was liberated from the slide and single-indexed libraries prepared using a 10x Genomics PCR-based protocol. Libraries were sequenced (1 per lane on a HiSeq4000), aiming for 300M raw reads per sample, with the following sequencing format; read 1: 28 cycles, i7 index: 8 cycles, i5 index: 0 cycles and read 2: 91 cycles.

### Alignment and quantification of scRNA-seq and snRNA-seq data

For each sequenced single-cell and single-nucleus RNA-seq library, we performed read alignment to the 10X Genomics’ GRCh38 3.0.0 human reference genome, mRNA version for scRNA-seq samples and pre-mRNA version for snRNA-seq samples, latter created following instructions from 10X Genomics: https://support.10xgenomics.com/single-cell-gene-expression/software/pipelines/latest/advanced/references#premrna. Quantification and initial quality control (QC) were performed using the Cell Ranger Software (version 3.0.2; 10X Genomics) using default parameters. Cell Ranger filtered count matrices were used for downstream analysis.

### Alignment and quantification of multiome data

For each sequenced snRNA-ATAC-seq (multiome) library, we performed read alignment to custom made genome consisting of 10X Genomics’ GRCh38 3.0.0 pre-mRNA human reference genome and 10X Genomics Cell Ranger-Arc 1.0.1 ATAC genome, created following instructions from 10X Genomics: https://support.10xgenomics.com/single-cell-multiome-atac-gex/software/pipelines/latest/advanced/references. Quantification and initial quality control (QC) were performed using the Cell Ranger-Arc Software (version 1.0.1; 10X Genomics) using default parameters. Cell Ranger-Arc filtered count matrices were used for downstream analysis.

### Downstream scRNA-seq and snRNA-seq analysis

#### Detection of doublets by gene expression

We used Scrublet for cell doublet calling on a per-library basis. We used a two-step diffusion doublet identification followed by Bonferroni-FDR correction and a significance threshold of 0.01, as described in^63^. Predicted doublets were not excluded from the initial analysis, but used afterwards to flag clusters with high doublet scores.

#### Detection of doublets by genotype

Souporcell ^64^ was used to deconvolute (a) maternal and fetal origin of cells and nuclei in our scRNA-seq and snRNA-seq samples (including multiome snRNA-seq); (b) assignment of cells to individuals in pooled samples (namely, samples Pla_HDBR8768477, Pla_HDBR8715512 and Pla_HDBR8715514); and (c) organoids from multiple individuals. In some samples deconvolution into maternal or fetal origin by genotype was not possible which is likely due to the highly skewed ratio of genotypes (either extremely high (>0.95) or extremely low (<0.05) ratio of maternal to fetal droplets). In those cases, maternal-fetal origin of the cells was identified using known markers from ^14^.

Souporcell (version 2.4.0) was installed as per instructions in https://github.com/wheaton5/souporcell and used in the following way:

path_to/singularity exec./souporcell.sif souporcell_pipeline.py -i
./cellranger_path/possorted_genome_bam.bam -b
./cellranger_path/filtered_feature_bc_matrix/barcodes.tsv -f./genome_path/genome.fa -t 8 -o souporcell_result -k 2 --skip_remap True --common_variants
./filtered_2p_1kgenomes_GRCh38.vcf

Where k=2 corresponds to the number of individuals to be deconvoluted (in our case either mother and fetus or pooled individuals H7 and H9 in samples Pla_HDBR8768477, Pla_HDBR8715512 and Pla_HDBR8715514. Accuracy of deconvolution was evaluated in downstream analysis once cluster identity was clear from either gene expression or predictions of logistic regression. In samples where deconvolution worked successfully, inter-individual doublets were further excluded from downstream analysis.

#### Filtering genes high in ambient RNA signal

To assess which genes in the scRNA-seq and snRNA-seq data were high in ambient RNA (soup) signal (further referred to as noisy genes), the following approach was undertaken separately for all the scRNA-seq and snRNA-seq samples:

1. Read in all the raw and filtered count matrices for each sample produced by Cell Ranger Software
2. Discard droplets with < 5 UMIs (likely to be fake droplets from sequencing errors)
3. Only keep data from samples which we further consider as noisy (where “Fraction reads in cells” reported by Cell Ranger is less than 70% (guided by 10X Genomics’ recommendations: https://assets.ctfassets.net/an68im79xiti/163qWiQBTVi2YLbskJphQX/e90bb82151b1cdab6d7e9b6c845e6130/CG000329_TechnicalNote_InterpretingCellRangerWebSummaryFiles_RevA.pdf)
4. Take the droplets that are in raw but are not in filtered matrices considering them as empty droplets
5. Concatenate all raw objects with empty droplets into 1 joint raw object and do the same for filtered
6. For all genes calculate soup probability as defined with the following equation:

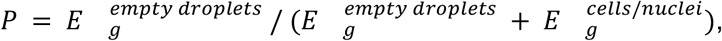

Where 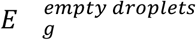 is the total sum of expression (number of UMI counts) of gene g in empty droplets, and, 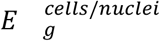 is the total sum of expression counts of gene g in droplets that are considered as cells/nuclei by Cell Ranger.
7. For all genes calculate number of cells/nuclei where the gene is detected at >0 expression level (UMI counts)
8. Label genes as noisy if their soup probability exceeds 50% quantile of soup probability distribution - done separately for cells and for nuclei

This approach was used to estimate noisy genes in (a) donor P13 samples and (b) all donors’ samples. Donor P13 noisy genes were excluded during mapping onto space (Visium, see section “Location of cell types in Visium data” below), whereas all donors’ noisy genes (labelled using nuclei-only derived threshold in step 8 to not over-filter genes based on the higher quality portion of the data which in our case in scRNA-seq) were excluded during all donors analysis of the whole atlas of all the cell states at the maternal-fetal interface.

#### Quality filters, alignment of data across different batches, and clustering

We integrated the filtered count matrices from Cell Ranger and analysed them with scanpy (version 1.7.1), with the pipeline following their recommended standard practises. Briefly, we excluded genes expressed by less than three cells, excluded cells expressing fewer than 200 genes, and cells with more than 20% mitochondrial content. After converting the expression space to log(CPM/100 + 1), the object was transposed to gene space to identify cell cycling genes in a data-driven manner, as described in ^63,65^. After performing PCA, neighbour identification and louvain clustering, the members of the gene cluster including known cycling genes (*CDK1, MKI67, CCNB2* and *PCNA*) were flagged as the data-derived cell cycling genes, and discarded in each downstream analysis where applicable.

Next, to have an estimate of the optimal number of latent variables to be used later in the single-cell Variational Inference (scVI) workflow for dimensionality reduction and batch correction, we identified highly variable genes, scaled the data and calculated PCA to observe the variance ratio plot and decide on an elbow point which defined values of n_latent parameter which were then used to correct for batch effect by 10X library batch (“sample”) with scVI. Number of layers in scVI models was tuned manually to allow for better integration. The resulting latent representation of the data was used for calculating neighbourhood graph, Uniform Manifold Approximation and Projection (UMAP) and further doing Louvain clustering. Analysis was done separately for (a) donor P13 trophoblast compartment and (b) all donors’ data (all cell states). In both analyses (a) and (b) trophoblast data was analysed separately with consecutive rounds of re-analysis upon exclusion of clusters of noisy nature (exhibiting gene expression characteristic of more than 1 distinct population). In addition, in all donors’ analysis fibroblast (maternal and fetal separately) and maternal NK, T, myeloid, epithelial, endothelial and perivascular compartments were re-analysed separately using the approach described in the previous paragraph to achieve fine grain annotation.

### Differential gene expression analysis

Differential gene expression analysis was performed with limma (limma version 3.46.0, edgeR version 3.32.1) with “cell_or_nucleus” covariate (scRNA-seq or snRNA-seq (including multiome snRNA-seq) origin of each droplet) backwards along the trajectory that was derived using stOrder approach, namely for the following 6 comparisons: VCT_CCC vs VCT (VCT and VCT-p cell states together); EVT-1 vs VCT_CCC; EVT-2 vs EVT-1; iEVT vs EVT-2; GC vs iEVT; eEVT vs EVT-2.

### Alignment, quantification, and quality control of multiome ATAC data

We processed scATAC-seq libraries coming from multiome samples (read filtering, alignment, barcode counting, and cell calling) with 10X Genomics Cell Ranger-Arc (version 1.0.1) using the pre-built 10X’s GRCh38 genome (version corresponding to Cellranger-arc 1.0.1) as reference. We called the peaks using an in-house implementation of the approach described in Cusanovich et al.^66^ (available at https://github.com/cellgeni/cellatac, revision 21-099). In short, the genome was broken into 5 kb windows and then each cell barcode was scored for insertions in each window, generating a binary matrix of windows by cells. Matrices from all samples were concatenated into a unified matrix, which was filtered to retain only the top 200K most commonly used windows per sample. Using Signac (https://satijalab.org/signac/ version 0.2.5), the binary matrix was normalised with term frequency-inverse document frequency (TF-IDF) followed by a dimensionality reduction step using Singular Value Decomposition (SVD). The first latent semantic indexing (LSI) component was ignored as it usually correlates with sequencing depth (technical variation) rather than a biological variation ^66^. The 2-30 top remaining components were used to perform graph-based Louvain clustering. Next, peaks were called separately on each cluster using macs2 ^67^. Finally, peaks from all clusters were merged into a master peak set (i.e. peaks overlapping in at least one base pair were aggregated) and used to generate a binary peak by cell-matrix, indicating any reads occurring in each peak for each cell.

This analysis was done separately for (a) all multiome data at first and (b) trophoblast only subset of the multiome data. In the latter analysis we used annotation labels from the RNA counterpart of the multiome samples to perform peak calling.

### Alignment, quantification, and quality control of Visium data

For each 10X Genomics Visium sample, we used Space Ranger Software Suite (version 1.1.0) to align to the GRCh38 human reference pre-mRNA genome (official Cell Ranger reference, version 3.0.0) and quantify gene counts. Spots were automatically aligned to the paired H&E images by Space Ranger software. All spots under tissue detected by Space Ranger were included in downstream analysis.

### Downstream analysis of 10X Genomics Visium data

#### Location of cell types in Visium data

To locate the cell states in the Visium transcriptomics slides, we used the cell2location tool v0.06-alpha ^68^. As reference, we used snRNA-seq data of donor P13. We used general cell state annotations from the joint all donors’ analysis (corresponding to donor P13 data), with the exception of the trophoblast lineage. Trophoblast annotations were taken from donor P13-only analysis of the trophoblast compartment. Using information about which genes are noisy (high in ambient RNA signal) in donor P13 snRNA-seq data (please see details in “Filtering genes high in ambient RNA signal” section above), we excluded those from the reference and Visium objects prior to cell2location model training which significantly improved the results of mapping (namely, eliminated off-target mapping of cell states, i. e. made results of mapping more specific to the correct anatomical regions). Following the tutorial: https://cell2location.readthedocs.io/en/latest/notebooks/cell2location_tutorial.html#Cell2location:-spatial-mapping, we trained cell2location model with default parameters using 10X library as a batch covariate in the step of estimation of reference cell type signatures. Results were visualised with scanpy (version 1.7.1). Plots represent estimated abundance of cell types (cell densities) in Visium spots.

#### Subsetting Visium data into anatomical regions with SpatialDE2

We used SpatialDE2 ^69^ tissue segmentation algorithm to assign Visium spots to three anatomical regions: (a) placenta; (b) decidua_and_villi_tips and (c) myometrium. We used mRNA abundances from the deconvolution results obtained with cell2location ^24^ in SpatialDE2 tissue segmentation. Assignment of obtained Visium spot clusters to regions was done upon visual inspection. Locations of certain fibroblast cell states indicative of the specific anatomical region (uterine smooth muscle cells, uSMC, and decidual stromal cells, dS, cell states) were also used to guide this assignment. In addition, low quality spots were discarded based on i) not being under tissue and, ii) having low count and gene coverage (visual inspection).

For more details, please refer to the following notebook: https://github.com/ventolab/MFI/blob/main/2_inv_troph_trajectory_and_TFs/2-1_stOrder_inv_troph/S1_regions_analysis_for_SpCov_model_and_later_for_CellPhone.ipynb

### Downstream snATAC-seq analysis

#### Quality filters

To obtain a set of high quality peaks for downstream analysis, we filtered out peaks that (i) were included in the ENCODE blacklist, (ii) have a width outside the 210-1500bp range and (iii) were accessible in less than 5% of cells from a *cellatac* cluster. Low quality cells were also removed by setting to 4 the minimum threshold for log1p transformed total counts per cell.

#### Alignment of data across different batches and clustering

We adopted the cisTopic approach^70,71^ for the core of our downstream analysis. cisTopic employs Latent Dirichlet Allocation (LDA) to estimate the probability of a region belonging to a regulatory topic (region-topic distribution) and the contribution of a topic within each cell (topic-cell distribution). The topic-cell matrix was used for constructing the neighbourhood graph, computing UMAP projections and clustering with the Louvain algorithm. After this was done for all cell states, clusters corresponding to trophoblast cell states (based on the unbiased clustering done here and annotation labels coming from the RNA counterpart of this multiome data) were further subsetted and re-analysed following the same pipeline.

#### Gene activity scores

Next, we generated a denoised accessibility matrix (predictive distribution) by multiplying the topic-cell and region-topic distribution and used it to calculate gene activity scores. To be able to integrate them with sc/snRNA-seq data, gene activity scores were rounded and multiplied by a factor of 10^7, as previously described ^71^.

#### Cell type annotation of invading trophoblast

Final labels of invading trophoblast in snATAC-seq data were directly transferred from RNA counterpart of the multiome data.

### StOrder: join inference of trophoblast invasion from gene expression and spatial data

StOrder is a computational framework for joint inference of cellular differentiation trajectories from gene expression data and information about location of cell states in physical space (further referred to as spatial data).

It consists of three principal steps:

1. Calculate pairwise cell state connectivity from gene expression data (here we use snRNA-seq data).
2. Calculate pairwise cell state proximity in physical space from spatial data (here we use Visium spatial transcriptomics data) using a new spatial covariance model.
3. Combine connectivity matrices from steps 1 and 2 in a weighted sum to reconstruct the putative tree structure of the differentiation trajectory.

First, StOrder relies on a gene expression-based connectivity matrix (generated in our case by PAGA ^72^) that establishes potential connections between cell state clusters defined by single cell/nucleus transcriptomics datasets. The values in this matrix can be interpreted as pairwise similarity scores for cell states in gene expression space. In our case we used snRNA-seq data from P13 as it contains all trophoblast subsets.

Second, StOrder generates a spatial covariance matrix that reflects pairwise proximity of cell states that co-exist in space and smoothly transition from one state to another while physically migrating in space. To do so, StOrder takes as an input the deconvolution results (derived in our case with cell2location ^24^) of Visium spatial transcriptomics data. Here, we used all spatial transcriptomics data profiled (donors P13, P14 and Hrv43). Then, it fits a Gaussian Process (GP) model that derives pairwise spatial covariance scores for all the cell state pairs with the following model:

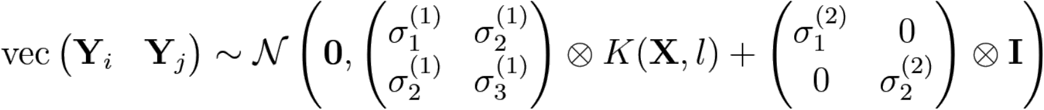

where 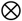 is the Kronecker product and the combined vector of cell densities **(Y_i,k_ Y_j,k_)** of cell states **i** and **j** is modelled by a multivariate Gaussian distribution whose covariance decomposes into a spatial and a noise term. The spatial term

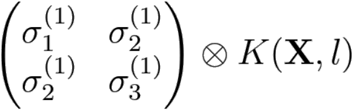

is defined by a between-cell-state covariance matrix

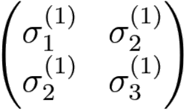

and a spatial covariance matrix *K*(X, *l*) defined using the squared exponential kernel:

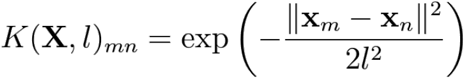

x_m_ and x_n_ are spatial coordinates of spots m and n and l is the length scale of the smooth GP function in space that is being fit to cell densities.

The noise term

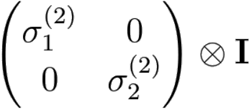

represents sources of variation other than spatial covariance of cell state densities.

The between-cell-state covariance matrix is constrained to be symmetric positive definite by defining

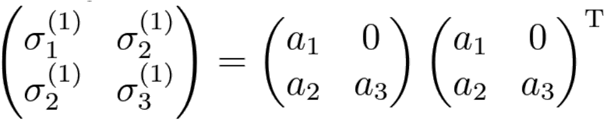

The free parameters a_1_, a_2_, a_3_, σ_1_^(2)^, σ_2_^(2)^, and | are estimated using maximum likelihood and automatic differentiation in Tensorflow ^73,74^ using the BFGS algorithm. To improve convergence, we initialise l to the distance between centres of neighboring Visium spots.

This model allows us to infer which cell states are proximal in physical space and are likely to be migrating in the process of gradual differentiation in space.

For the spatial covariance model within StOrder workflow we only used a subset of our Visium data that corresponded to (a) decidua_and_villi_tips and (b) myometrium - because only these regions contained invading trophoblast cell states. For more details please see section “Subsetting Visium data into anatomical regions with SpatialDE2” in “Downstream analysis of 10x Genomics Visium data” above. This helps to focus on the regions of the tissue that are relevant for the process of interest and is recommended to do in general if there are parts of the Visium data that do not contain cell states relevant to the process of interest.

Third, StOrder reconstructs connections between cell states by taking into account both the connectivity matrix (step 1) from single-cell transcriptomics data and the spatial covariance matrix (step 2) from the spatial data by summing the two matrices in a weighted manner and reconstructing the putative trajectory tree using the built-in PAGA functions.

The combined connectivity matrix based on both gene expression and spatial data with a range of weight parameters (0.16 ≤ω≤ 0.47 for gene expression weight/contribution) revealed the fully resolved invasion trajectory tree of the EVT with the correct topology (all connected cell state components, one branching point, no cycles, start at VCT-CCC population and two endpoints: eEVT and GC populations). The choice of ω parameter (contribution/weight of gene expression vs spatial part in the final matrix) in this last step depends on the goal of using this approach. In our case, we assumed: (i) the origin of EVT (VCT-CCC); (ii) the endpoints of EVT (eEVT and GC); (iii) the determination of a single branching point; and (iv) the absence of cyclic trajectory. We therefore produced trajectory trees for 101 values of ω parameter (from 0 to 1 with 0.01 increment step) representative of different tree topologies corresponding to different ratios of gene expression vs spatial contribution. Out of the 101 tree structures we inspected for ω values in the [0.16, 0.47] interval the trees represented the topology with the assumptions described above. These trajectories consistently assigned EVT-2 as the putative branching point. Tree structures for ω > 0.47 (mainly gene expression based connectivities) values did not yield a branching point population we were looking for. Tree structures with ω < 0.16 (mainly spatial based connectivities) hindered the link between iEVT and GC populations, likely due to the large length scale of this invasion in space.

#### Limitations

Our approach assumes the gradual nature of gene expression changes accompanied by gradual migration of cells in space while they differentiate. Thus, it may not yield meaningful results in scenarios where this underlying assumption is violated. In addition, it is recommended that the user estimates the spatial scale at which the process of interest is taking place - whether in current Visium resolution the differentiation and migration is happening over the course of only a few spots or many more - this will change the initial values of l parameter and help the model fit thedata better.

### Combined RNA/ATAC analysis using MEFISTO

#### Preprocessing of multiome data and training of the MEFISTO model

Gene expression (snRNA-seq) counts of the multiome data for donor P13 were normalised by total counts (scanpy.pp.normalize_per_cell(rna, counts_per_cell_after=1e4)) and log transformed (pp.log1p(rna)). Highly variable gene features were then calculated (sc.pp.highly_variable_genes(rna, min_mean=0.0125, max_mean=3, min_disp=0.5)) and the subsetted object’s expression was scaled (sc.pp.scale(rna, max_value=10)).

Chromatin accessibility (scATAC-seq) counts of the multiome data for donor P13 were preprocessed using TF-IDF normalisation (muon.atac.pp.tfidf(atac[key], scale_factor=1e4)). To select biologically meaningful highly variable peak features, ATAC counts were aggregated into pseodubulks by cell states and averaged, then variance of accessibility was calculated across these pseudobulks, and informative peak features were selected based on this measure (top 75th percentile (10640) of peaks selected in total) as the peaks with highest variance. Lastly, this data was scaled (sc.pp.scale(atac, max_value=10)).

Using the preprocessed RNA and ATAC data we used a pseudotime-aware dimensionality reduction method MEFISTO^37^ to extract major sources of variation from the RNA and ATAC data jointly and identify coordinated patterns along the invasion trajectory. As a proxy for the trophoblast invasion trajectory in the MEFISTO model we used 2-dimensional pseudotime coordinates based on a UMAP of the RNA data by calculating PCA (sc.tl.pca(rna, n_comps=8)), neighborhood graph (sc.pp.neighbors(rna)) and UMAP embedding (sc.tl.umap(rna)).

The MEFISTO model was trained using the following command within MUON (version 0.1.2) package interface:

~~~
muon.tl.mofa(mdata, outfile=”,
  use_obs = “union”,
  smooth_covariate=[“UMAP1”, “UMAP2”],
  use_float32=True)
~~~

We further excluded factor 5 from downstream analysis as a technical artefact due to its significant and high correlation (Spearman rank-order correlation coefficient 0.94 (over all cell states), p-value < 10e-308, two-sided test) with the n_peaks_by_counts (number of ATAC peaks with at least 1 count in a nucleus) in ATAC view in all cell states (**Supp. Fig. 4k**) and lack of smoothness along pseudotime (**Supp. Fig. 4j**).

#### Defining groups of ATAC peak features

To further interpret ATAC features, we annotated them based on their genomic location using GenomicRanges package (version 1.42.0). In parallel, we used epigenetic data from^75^ to mark peak features in close proximity to trophoblast-specific enhancer features. To do so, we used peak files corresponding to H3K4me1, H3K27ac and H3K27me3 histone modifications marks for second trimester trophoblast samples (obtained from authors of aforementioned study upon request) to infer regions of the genome corresponding to active (H3K27ac + H3K27me3), primed (only H3K4me1) or repressed (H3K4me1 + H3K27me3) enhancers. This was done using bedtools (version 2.30.0) in the following way:

1. bedtools subtract -a H3K4me1_file.bed -b H3K27ac_file.bed > interm_file.bed bedtools subtract -a interm_file.bed -b H3K27me3_file.bed > primed_enhancers.bed To produce primed enhancers file
2. bedtools intersect -a H3K4me1_file.bed -b H3K27ac_file.bed > active_enhancers.bed To produce active enhancers file
3. bedtools intersect -a H3K4me1_file.bed -b H3K27me3_file.bed > repressed_enhancers.bed To produce repressed enhancers file

The enhancer files produced were then overlapped with peaks in ATAC analysis (bedtools intersect -a atac_peaks_file.bed -b enhancer_file.bed -wa) and any peaks having a >1bp overlap with an enhancer feature were considered to be proximal to those features (done separately for active, primed and repressed enhancers).

#### Enrichment analysis of features in the MEFISTO model

Gene set enrichment analysis for gene features was performed based on the C5 category and the Biological Process subcategory from the MSigDB database (https://www.gsea-msigdb.org/gsea/msigdb) using GSEA functionality implemented in MOFA2 (run_enrichment command, MOFA2 version 1.3.5). This was done separately for negative and positive weights of each factor.

Peak group enrichment for peak features was performed using the same run_enrichment command in MOFA2 on peak groups defined as described above (Defining groups of ATAC peak features).

#### TF analysis using the MEFISTO model

To extract information about TF binding motif enrichment in ATAC features of MEFISTO factors, we first performed enrichment analysis of peaks using GSEA functionality implemented in MOFA2 (run_enrichment command, MOFA2 version 1.3.5) on the peak-motif matrix produced by Signac package (version 1.5.0). Then, to identify which MEFISTO factors contribute the most to each transition of cell states along the invading trophoblast trajectory (inferred with StOrder), we trained logistic regression classifiers for each transition along the trajectory (overall for 6 transitions: VCT → VCT-CCC, VCT-CCC → EVT-1, EVT-1 → EVT-2, EVT-2 → iEVT, iEVT → GC, EVT-2 → eEVT) on the matrix of factor values. For each transition the factor with the highest absolute coefficient separating the two cell states was selected, accounting for the sign of contribution in the logistic regression (positive or negative). If the top factor is contributing to a transition with a positive coefficient, TF binding motifs coming from MEFISTO enrichment analysis of this factor’s top positive values are further considered in general TF analysis as TFs upregulated upon this transition, whereas TF binding motifs coming from MEFISTO enrichment analysis of this factor’s top negative values are further considered in general TF analysis as TFs downregulated upon this transition. All of these TF motifs are marked as having evidence from the MEFISTO factor relevant for this transition. Reverse procedure is applied in case if the top factor is contributing to a transition with a negative coefficient in the corresponding logistic regression model.

For more details please see the following notebook: https://github.com/ventolab/MFI/blob/main/2_inv_troph_trajectory_and_TFs/2-5_MEFISTO_analysis_inv_troph/S3_DEG_comparison_to_MEFISTO_factor_translation.ipy_nb

### CellPhoneDB and CellSign

To retrieve interactions between invading trophoblast and other cell populations identified in our samples, we used CellPhoneDB v4 ‘`degs_analysis`’ method ^14^’^76^(https://github.com/ventolab/CellphoneDB) described in ^23^. In short, we retrieved the interacting pairs of ligands and receptors meeting the following requirements: 1) all the protein members were expressed in at least 10% of the cell type under consideration; and 2) at least one of the protein members in the ligand or the receptor was a differentially expressed gene in an invading trophoblast subset (according to our analysis of differential expression, for details please see section “Differential gene expression analysis” above), with an adjusted p-value below 0.05. We further selected which cell states are spatially co-located in each microenvironment via visual inspection of cell2location deconvolution results for our Visium data.

### Transcription Factor (TF) analysis

To prioritise the TFs relevant for each invading trophoblast cell state or microenvironment, we integrate four types of measurements: (i) expression levels of the TF and (ii) the activity status of the TF measured from (ii-a) the expression levels of their targets (described below in *“Transcription factor activities derived from scRNA-seq and snRNA-seq”*) and/or (ii-b) the chromatin accessibility of their binding motifs (described below in *“Transcription factor motif activity analysis from scATACseq”*) and/or (ii-c) evidence of the chromatin accessibility of their binding motifs in relevant factors from multimodal RNA-ATAC analysis (with MEFISTO). Plots in main figures include TF meeting the following criteria: 1) TF was differentially expressed, with adjusted p-value < 0.01), and/or 2) TF was differentially active, with log2 fold change greater than 0.75 and adjusted p-value < 0.01 in at least one of the TF activity measurements (iia/iib).

#### Transcription factor differential expression (from scRNAseq and snRNA-seq)

We compute differential expression using the procedure described in section “Differential gene expression analysis” above and further subset resulting gene targets to TFs only based on the list of TFs provided by DoRothEA.

#### Transcription factor activities derived from scRNAseq and snRNAseq

We estimated protein-level activity for human Transcription factors (TF) as *a proxy* of the combined expression levels of their targets. Target genes were retrieved from *Dorothea ^77^*, an orthogonal collection of TF targets compiled from a range of different sources. Next, we estimated TF activities for each cell using *Viper^78^*, a GSEA-like approach, as implemented in the *Dorothea* R package and tutorial ^79^ for the genes differentially expressed along the invading trophoblast trajectory (see section “Differential gene expression analysis” above).

#### Transcription factor motif activity analysis from scATACseq

Transcription factor motif activities were computed using chromVar ^80^ v. 1.12.2 with positional weight matrices from JASPAR2018 ^81^, HOCOMOCOv10 ^82^, SwissRegulon ^83^, HOMER ^84^. chromVar returns a matrix with binding activity estimates of each TF in each cell, which we used to test for differential TF binding activity between trophoblast cell states with FindMarkers function in Seurat (default parameters) in the same way as described in section “Differential gene expression analysis” above (backwards along invading trophoblast trajectory).

