## Supplementary material for "Spatially resolved single-cell multiomics map of human trophoblast differentiation in early pregnancy": supllemental files

**Supplementary Table 8. Supplementary_Table8.xlsx (separate file)**
Probes used for multiplexed RNAscope smFISH.

Supplementary Figure 1

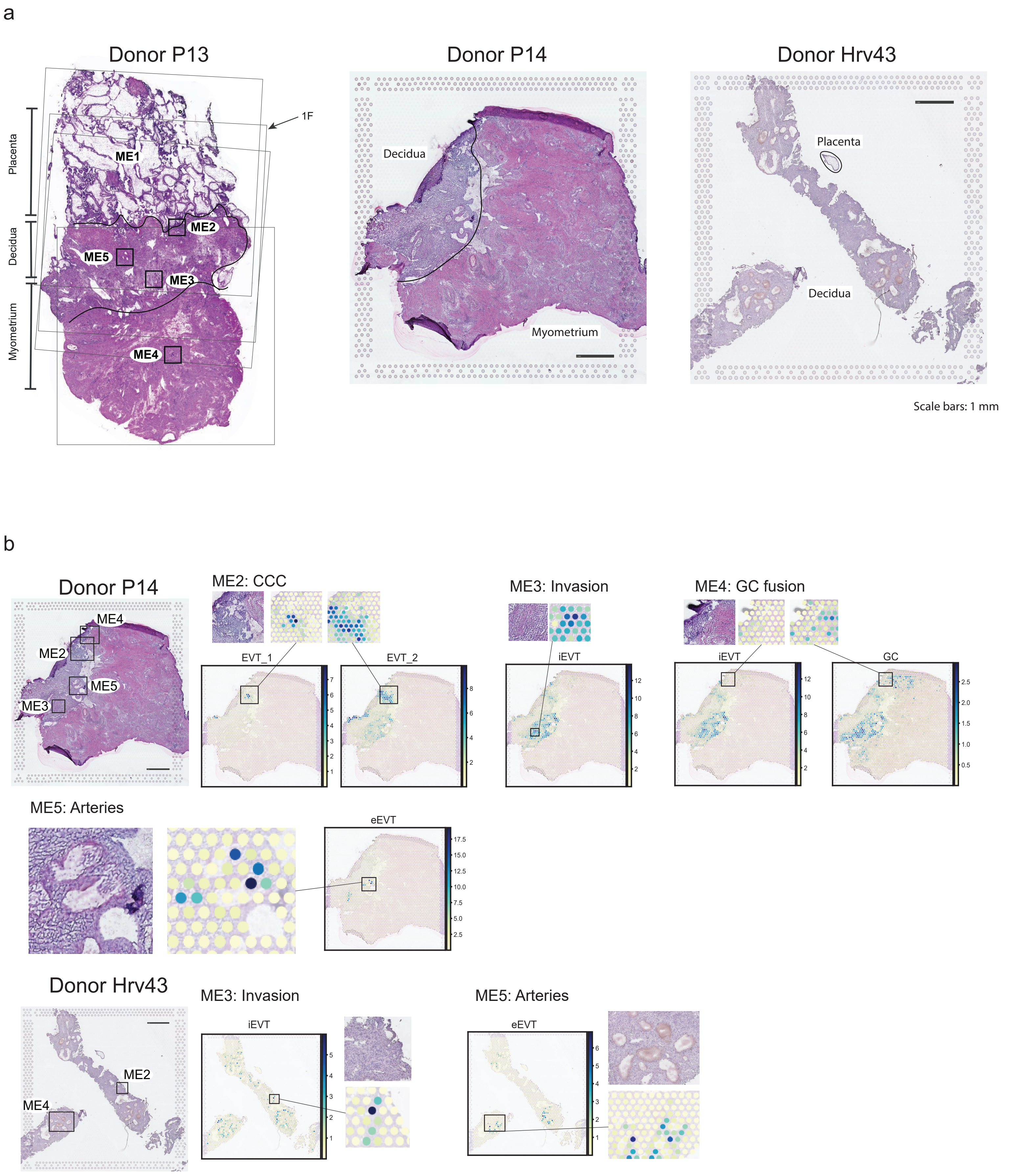

### Supplementary Figure 2

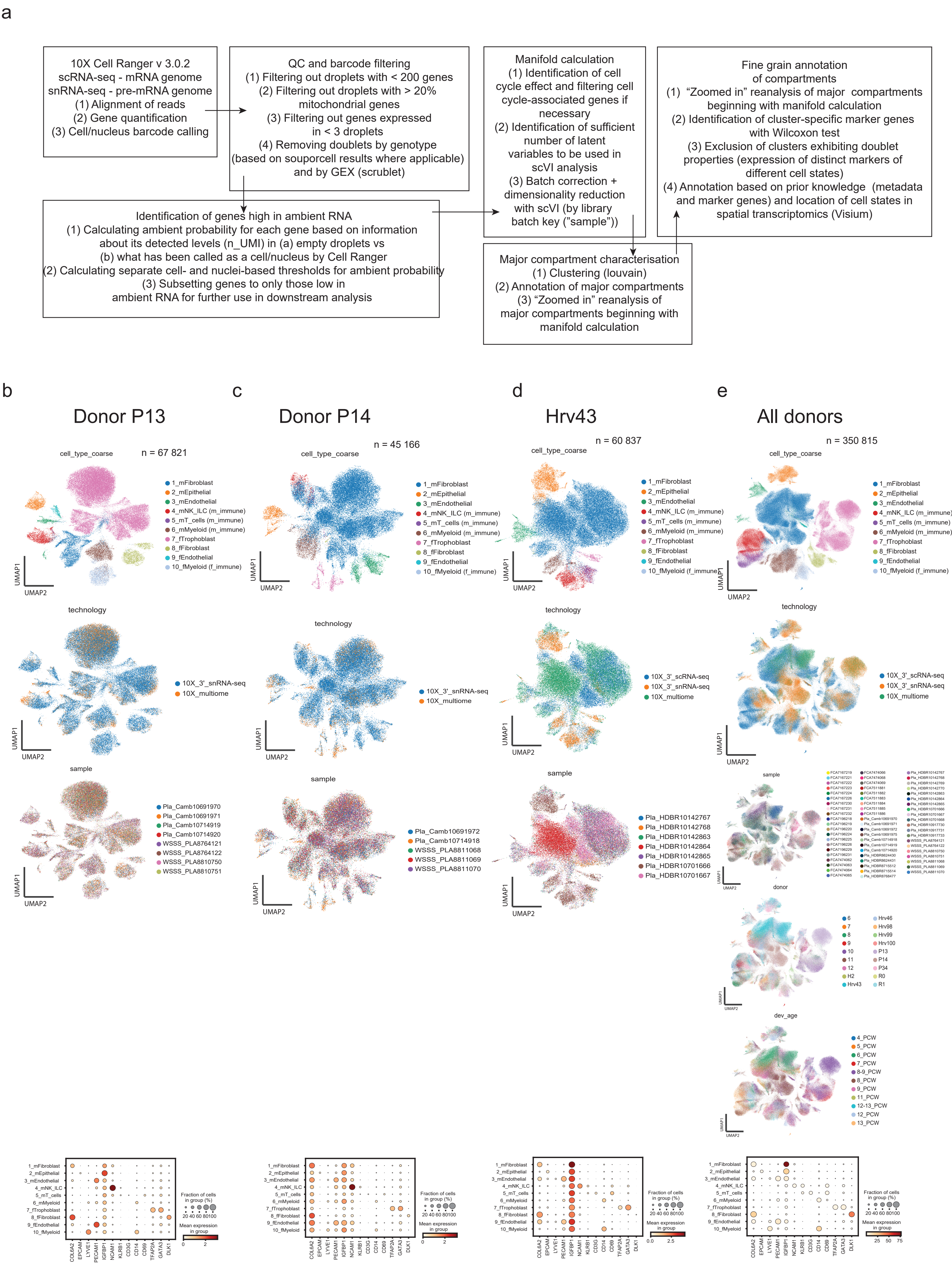

### Supplementary Figure 3

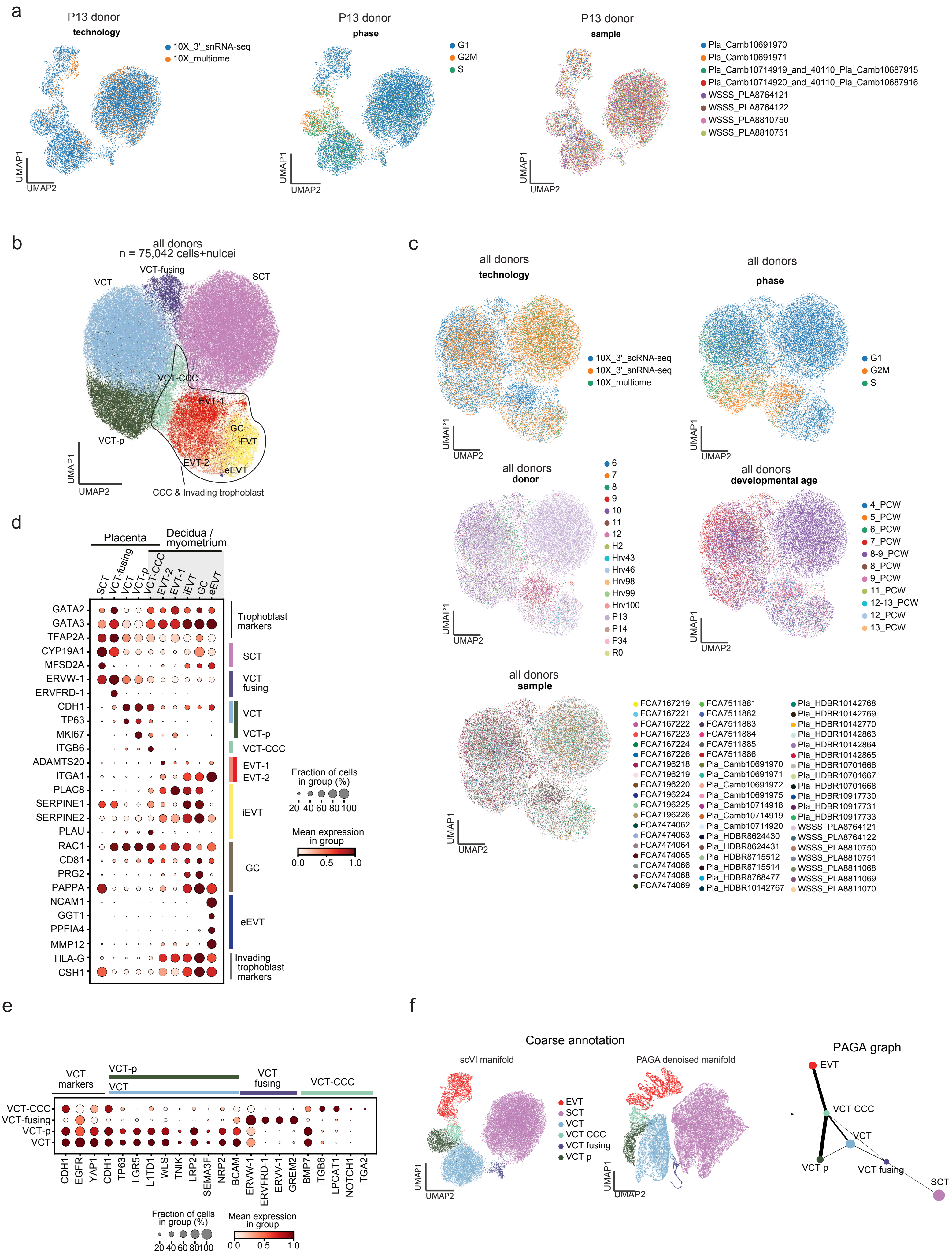

#### Supplementary Figure 4

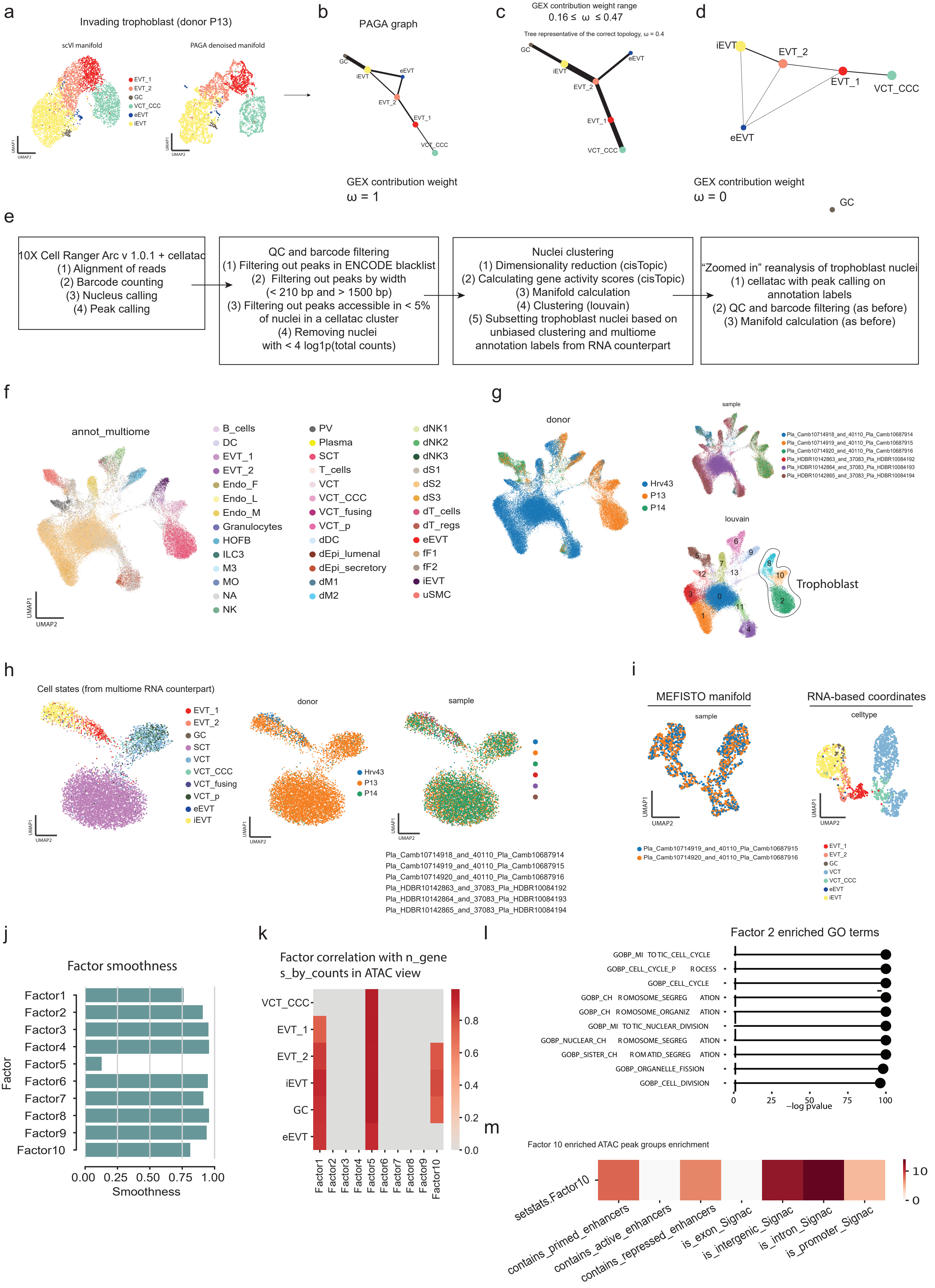

Supplementary Figure 5

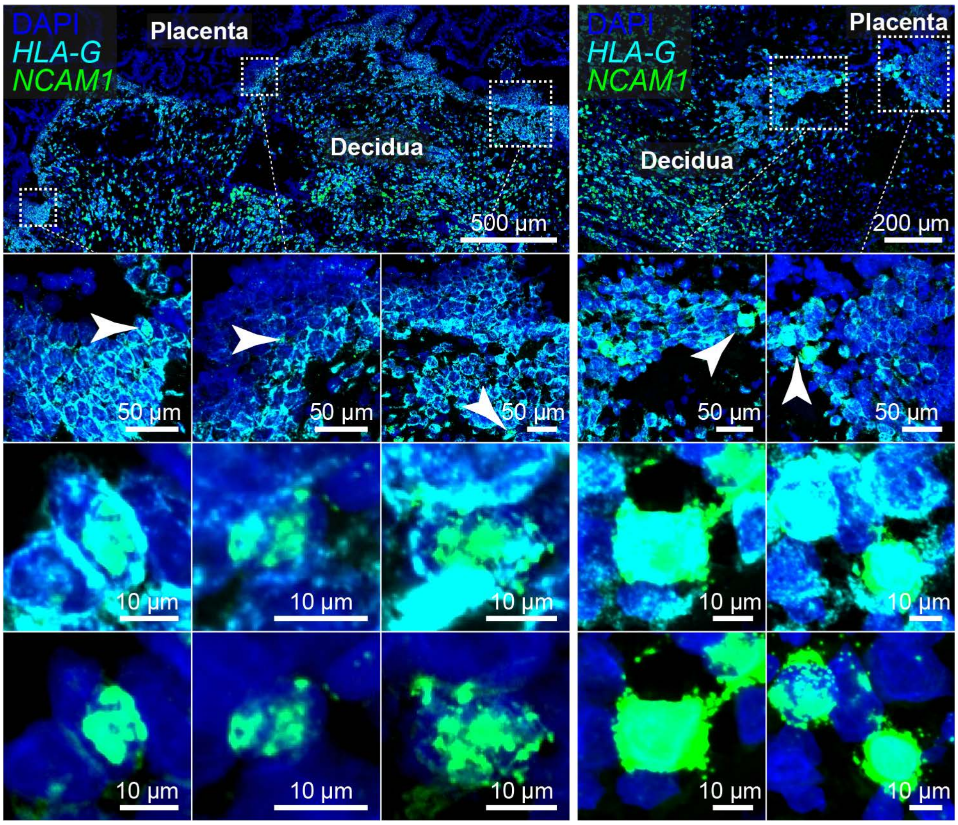

### Supplementary Figure 6

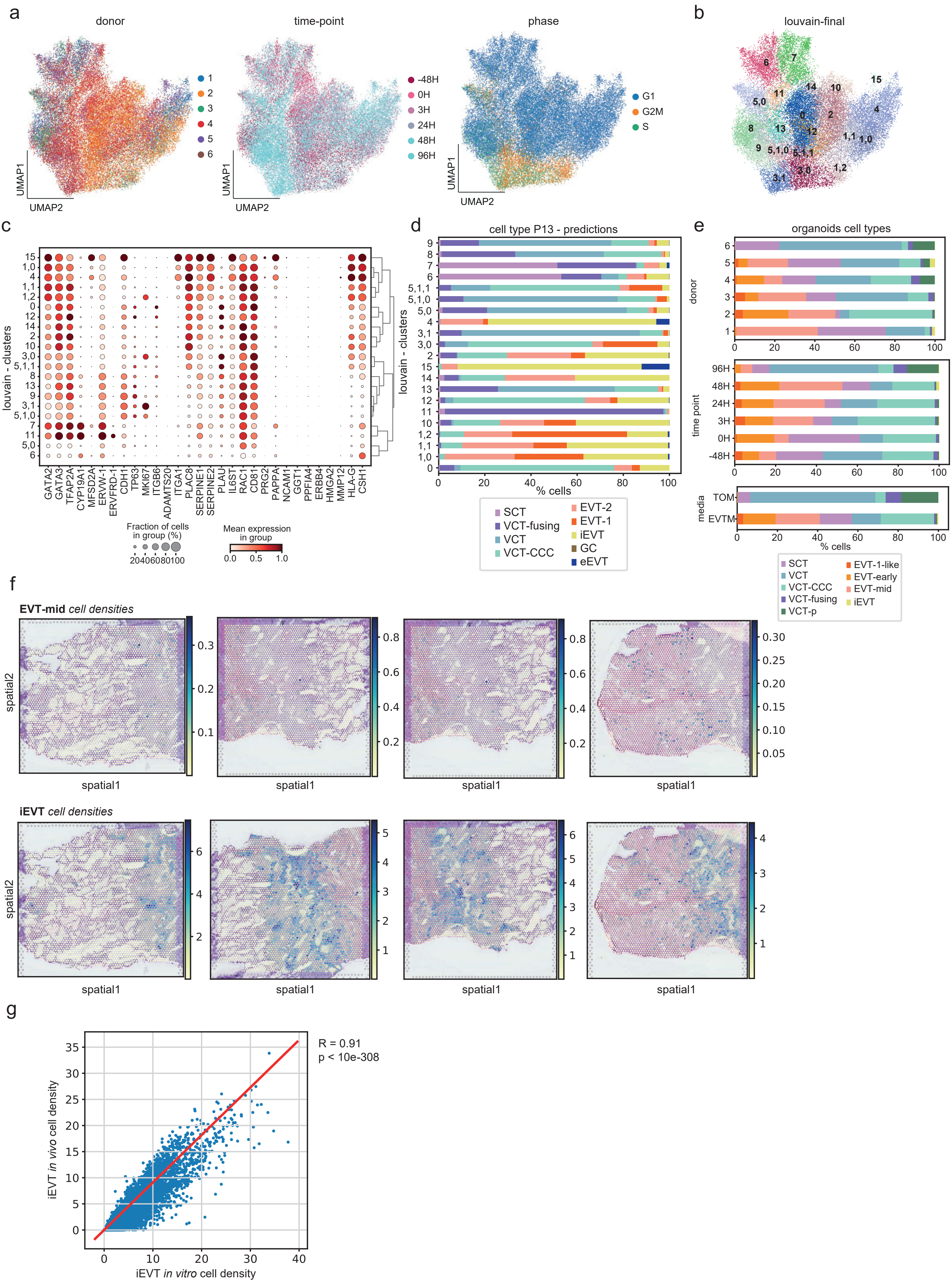
